# Similar but distinct: the impact of biomechanical forces and culture age on the production, cargo loading, and biological efficacy of human megakaryocytic extracellular vesicles for applications in cell and gene therapies

**DOI:** 10.1101/2023.04.26.538401

**Authors:** Will Thompson, Eleftherios Terry Papoutsakis

## Abstract

Megakaryocytic extracellular vesicles (MkEVs) promote the growth and megakaryopoiesis of hematopoietic stem and progenitor cells (HSPCs) largely through endogenous miR-486-5p and miR-22-3p cargo. Here, we examine the impact of biomechanical force and culture age/differentiation on the formation, properties, and biological efficacy of MkEVs. We applied biomechanical force to Mks using two methods: shake flask cultures and a syringe pump system. Force increased MkEV production in a magnitude-dependent manner, with similar trends emerging regardless of whether flow cytometry or nanoparticle tracking analysis (NTA) was used for MkEV counting. Both methods produced MkEVs that were relatively depleted of miR-486-5p and miR-22-3p cargo. However, while the shake flask-derived MkEVs were correspondingly less effective in promoting megakaryocytic differentiation of HSPCs, the syringe pump-derived MkEVs were more effective in doing so, suggesting the presence of unique, unidentified miRNA cargo components. Higher numbers of MkEVs were also produced by “older” Mk cultures, though miRNA cargo levels and MkEV bioactivity were unaffected by culture age. A reduction in MkEV production by Mks derived from late-differentiating HSPCs was also noted. Taken together, our results demonstrate that biomechanical force has an underappreciated and deeply influential role in MkEV biology, though that role may vary significantly depending on the nature of the force. Given the ubiquity of biomechanical force in vivo and in biomanufacturing, this phenomenon must be grappled with before MkEVs can attain clinical relevance.

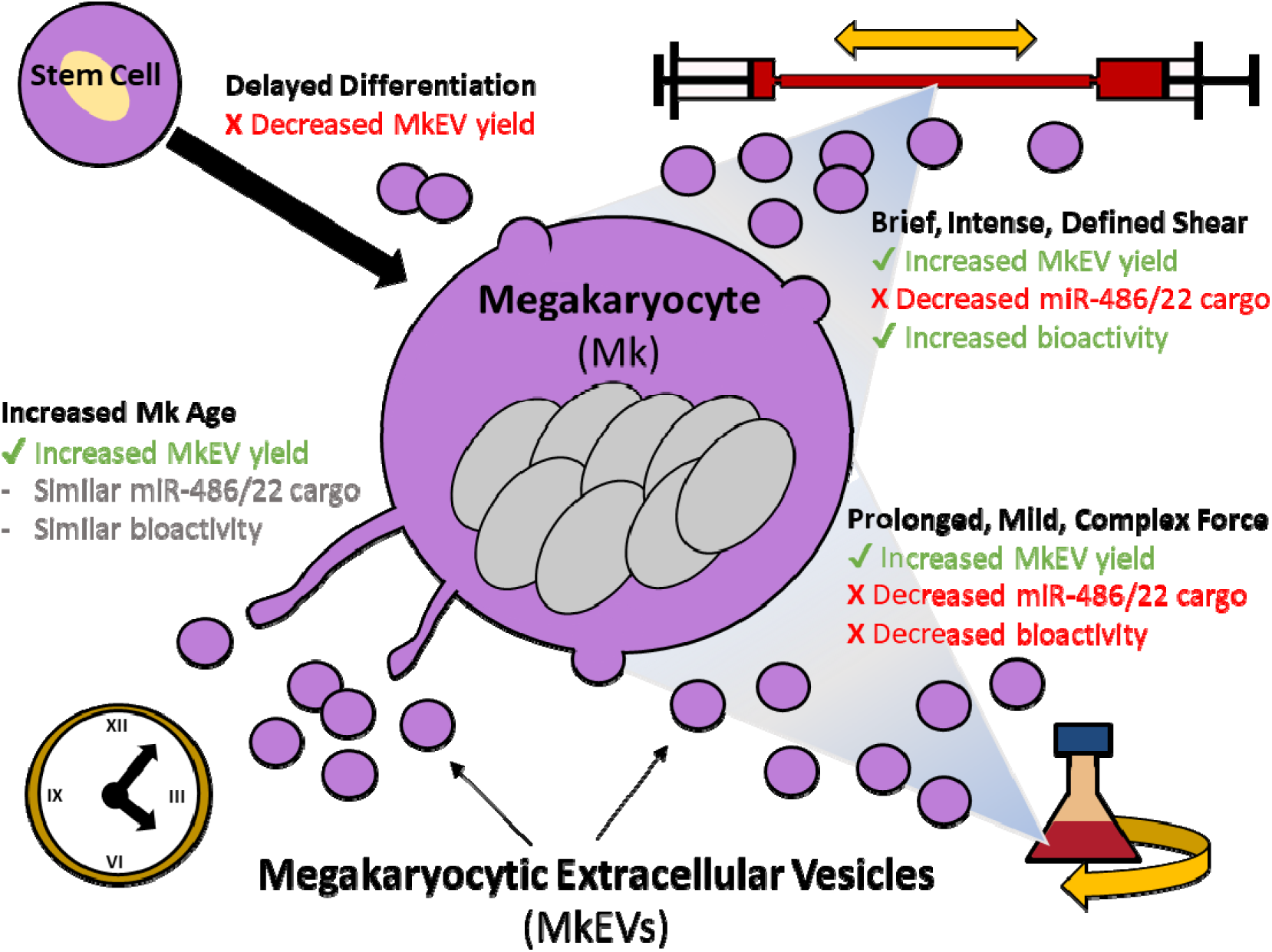

## 1 INTRODUCTION

Megakaryocytes (Mks) are large mammalian platelet-producing cells. Under exposure to a cocktail of appropriate cytokines—most notably thrombopoietin (TPO)—hematopoietic stem and progenitor cells (HSPCs) in the bone marrow differentiate to form Mks.^1–3^ As they mature, Mks become polyploid, accumulating nuclei without dividing, and produce branching extensions called “proplatelets” with which they burrow through the walls of nearby sinusoidal blood vessels in the bone marrow.^1^ Subjected to the shear force in these vessels, proplatelets fragment and detach from their parent Mks, forming platelet precursors known as preplatelets.^1,4^ More recently, research has also demonstrated the migration of whole Mks into the lung vasculature, suggesting a pulmonary origin for some platelets.^5^

Extracellular vesicles (EVs) are submicron-size particles with lipid bilayer membranes that are released by every cell type. They contain protein, lipid and nucleic acid cargo, which they deliver to target cells, often as a means of mediating cellular phenotype.^6–8^ Previously, EVs were categorized as either exosomes (Exos) or microparticles (MPs) on the basis of their biogenesis. MPs are produced directly from the outward budding of the plasma membrane, while Exos originate when exocytosis of multivesicular bodies (MVBs) results in the release of intraluminal vesicles—formed via the inward budding of MVB membranes—into the extracellular space.^6,7^ Exos, at <100 nm in diameter, are generally smaller than MPs (100-1,000 nm in diameter) and have membranes enriched in endosomal proteins. MPs are of special interest for scaled production due to their larger size—which enables the transport of extra cargo—and their relatively simpler isolation. Nevertheless, difficulty in separating the EV subtypes by biogenesis method has led the field to embrace the more general “EV” label.^9^ Therefore, we use the abbreviation “MkEVs” to describe the megakaryocytic EVs produced in this study, though our methods (i.e., ultracentrifugation at 25,000 × g for 30 min.) will concentrate larger particles (formerly MkMPs) and omit many smaller particles (formerly MkExos).

Through the combinatorial effects of their surface receptors, small RNA content, and other cargo, EVs have extensive applications in cell and gene therapies. EVs use various surface receptors to bind specifically to target cells, which subsequently internalize the bound EVs and/or their cargo by way of either endocytosis or membrane fusion.^10,11^ In many cases, the internalized cargo mediates cellular phenotype.^8,12^ For instance, MkEVs have been shown to uniquely target HSPCs, with tetraspanins CD54, CD11b, CD18, and CD43 acting as key mediators of MkEV-HSPC binding.^11^ Subsequent MkEV cargo delivery to the target HSPCs promotes megakaryopoiesis (differentiation into Mks), even in the absence of TPO.^13^ Our group has confirmed this function in vivo by using MkEVs to alleviate thrombocytopenia in murine models,^14^ a phenomenon which may explain recent reports that MkEV levels are lower in human patients with immune thrombocytopenia.^15^ Thus, MkEVs offer promise as a treatment for thrombocytopenia and other various Mk/platelet disorders, a finding which takes on particular importance given the severity of current platelet shortages.^16,17^ Even absent endogenous cargo, MkEVs can be loaded with synthetic cargo and utilized exclusively as delivery vehicles on the basis of their unique capacity for uptake by HSPCs.^18^

MicroRNAs (miRNAs) are small, single-stranded RNAs of ~22 nucleotides that influence cell phenotype by binding to—and subsequently silencing—messenger RNA (mRNA). Along with protein cargo, miRNA cargo is responsible for the action of EVs on target cells.^19^ For our system here, the megakaryopoiesis-promoting function of MkEVs can be almost entirely explained by the synergistic action of miR-486-5p and miR-22-3p cargo on HSPCs.^20^ These miRNAs may mediate JNK and PI3K/Akt/mTOR signaling, with miR-486-5p governing early megakaryopoiesis and miR-22-3p largely responsible for Mk maturation.^20^ Other work has suggested a role for miR-125b, miR-99a, and/or C-X-C chemokine receptor type 4 (CXCR4), all of which may activate Notch1 via the downregulation of DNA methyltransferases.^21^

EVs from a particular source have been generally treated as known quantities with unchanging properties. However, a great number of variables—including cell density, cell passage, oxygen and nutrient availability, temperature and pH stress, substrate quality, chemical and oxidative stress, irradiation, and electrical, mechanical, and acoustic stimulation—have been found to regulate EV cargo and function.^22–24^ We draw parallels with cellular protein glycosylation, which has only recently been appreciated as highly culture- and process-dependent.^25^

For our system here, while the specific binding (tropism) to HSPCs and megakaryopoiesis-promoting function of MkEVs is well-established both in vitro and in vivo, nothing is known about the potential variability of MkEV quality that occurs under different culture conditions. Mks are notoriously sensitive to mechanical stimulation, with shear and/or turbulence linked to both increased platelet release^26,27^ and faster aging/maturation.^13,28^ Higher (by up to 47-fold) MkEV yields have been observed following brief (0.5-2 h) exposure of mature Mks to shear stress,^13^ but the cargo and efficacy of these MkEV samples were never investigated. Understanding the correlations between MkEV quantity and quality is crucial, however, since low yields and unpredictably heterogenous samples currently pose the greatest impediments to large-scale EV production.^29^ Thus, we hypothesized that MkEV yields, structural characteristics, cargo, and bioactivity may be impacted by biomechanical force and other, related factors such as Mk and HSPC age. Our results demonstrate for the first time that dramatic differentials in MkEV quality occur in response to various magnitudes, durations, and types of biomechanical force on parent Mks. The impacts of Mk and HSPC age, however, are largely confined to variations in MkEV yield.

## 2 RESULTS

### 2.1 Experimental design and rationale

#### 2.1.1 The biomechanical force experiments

As noted in the Introduction, the Mk environment in vivo is far from static. In the sinusoids of the bone marrow, Mks are exposed to shear stress ranging from 1.3-4.1 dyn/cm^2,4^ while the vessel wall shear stress they encounter during migration to the lung vasculature averages 10-15 dyn/cm^2,30^ with possibility for a wide range of higher values.^31^ Given the magnitude of MkEV production required for clinical relevance, shear levels in Mk bioreactors will undoubtedly be higher still. Thus, we imposed two models of biomechanical force on parent Mks: long-term, complex rotational mixing via shake flasks, and brief, defined, and high-intensity shear stress in a novel syringe pump system.

The first model (rotational mixing via shake flasks) is more akin to the complexity of biomechanical force in industrial bioreactors, while the second model (syringe pump system) mimics the average wall shear stress (WSS) observed in the pulmonary vasculature and is easier to scale. For the shake flask experiments, Mks were transferred from T-flasks to shake flasks on day 10 of standard 12-day culture and rotated at either 60 or 120 rpm for two days. MkEVs were isolated on day 12 and subsequently analyzed according to the methods set forth below. Similarly, for the syringe pump experiments, 20 mL of Mks from standard day 12 culture were directly loaded into two syringes connected by 250 mm of a 1.58 mm ID silicone tube. The syringes were attached to separate syringe pumps and alternately discharged at a rate of 4,448 mL/h for 1.5 h, which corresponded to an average WSS of 15 dyn/cm^2^ in the connective tubing, where each Mk spent an average of 2.5% of its time. A Reynolds number of 1,000 within the connective tubing confirms the flow is laminar. Viability of the cells subjected to the syringe pump treatment was not significantly different from control cells (data not shown). We note that because the Mks were suspended in media (i.e., not adherent), they were exposed to a complex range of forces far beyond simple shear, including significant mechanical stress as a result of the rapid expansion/constriction of flow fields that occurred at the entrance of each syringe. Note that in all the aforementioned experiments, MkEVs created prior to the application of biomechanical force were included in the MkEV samples.

#### 2.1.2 The Mk culture age experiments

Given the established correlation between shear stress and Mk maturation, we hypothesized that Mk age may impact. MkEVs from normal (static) Mk cultures were collected after 11, 12 and 13 days and subsequently analyzed per the description later in this section.

#### 2.1.3 The “recycle” experiments examining HSPC age and passage

Given the large number (~80%) of non-Mks (i.e., CD41^−^/CD61^−^ cells) removed from D7 culture per our established protocols,^11,13^ we hypothesized that Mk yield could be increased by “recycling” these non-Mks through our culture process. With this in mind, CD41^−^/CD61^−^ cells removed from Mk (CD41^+^/CD61^+^) cultures on D7 were resuspended in D0 media and treated as D0 cells. This process was then repeated a second time for the second “generation” of D7 cultures. Since many or most of the cells in the CD41^−^/CD61^−^ fractions were CD34^+^ (Fig. 8B), re-culturing them is akin to “passaging” undifferentiated HSPCs and allows for an examination of the ways in which delayed HSPC differentiation impacts eventual Mk productivity (i.e., MkEV production).

### 2.2 Long-term, complex biomechanical forces in shake flasks enhance MkEV production in a force-magnitude dependent manner without affecting MkEV size

A schematic describing the shake flask experiments is present in Fig. 1A. MkEVs were counted via both flow cytometry (Fig. 1B) and NTA (Fig. 1C) and are plotted on a per-Mk basis. Both methods produced similar trends, though NTA-derived counts were roughly 2 orders of magnitude higher than those calculated using flow cytometry. This phenomenon, reflected elsewhere in the literature, is likely due to an undercount of small (<200 nm) or CD41^−^ EVs when using traditional flow cytometry and an overcount of non-EV particles (e.g., protein aggregates or lipoproteins) when using NTA.^32^ Nevertheless, both counting methods suggest that biomechanical force boosts MkEV production, and this trend further appears to be force-magnitude dependent, with progressively—and often significantly—higher MkEV counts observed with each increase in rotational speed (Fig. 1B-C). Using NTA to measure MkEV size distribution (Fig. 1D-F), we found that there were no significant differences in the mean diameter of the particles. Moreover, flow cytometry was employed to measure the percentages of MkEVs expressing CD54 and CD11b, two known mediators of MkEV-HSPC binding.^11^ Here again, no significant differences were observed, and CD11b levels in the MkEVs were low enough as to be considered negligible (data not shown).

**Figure 1.**
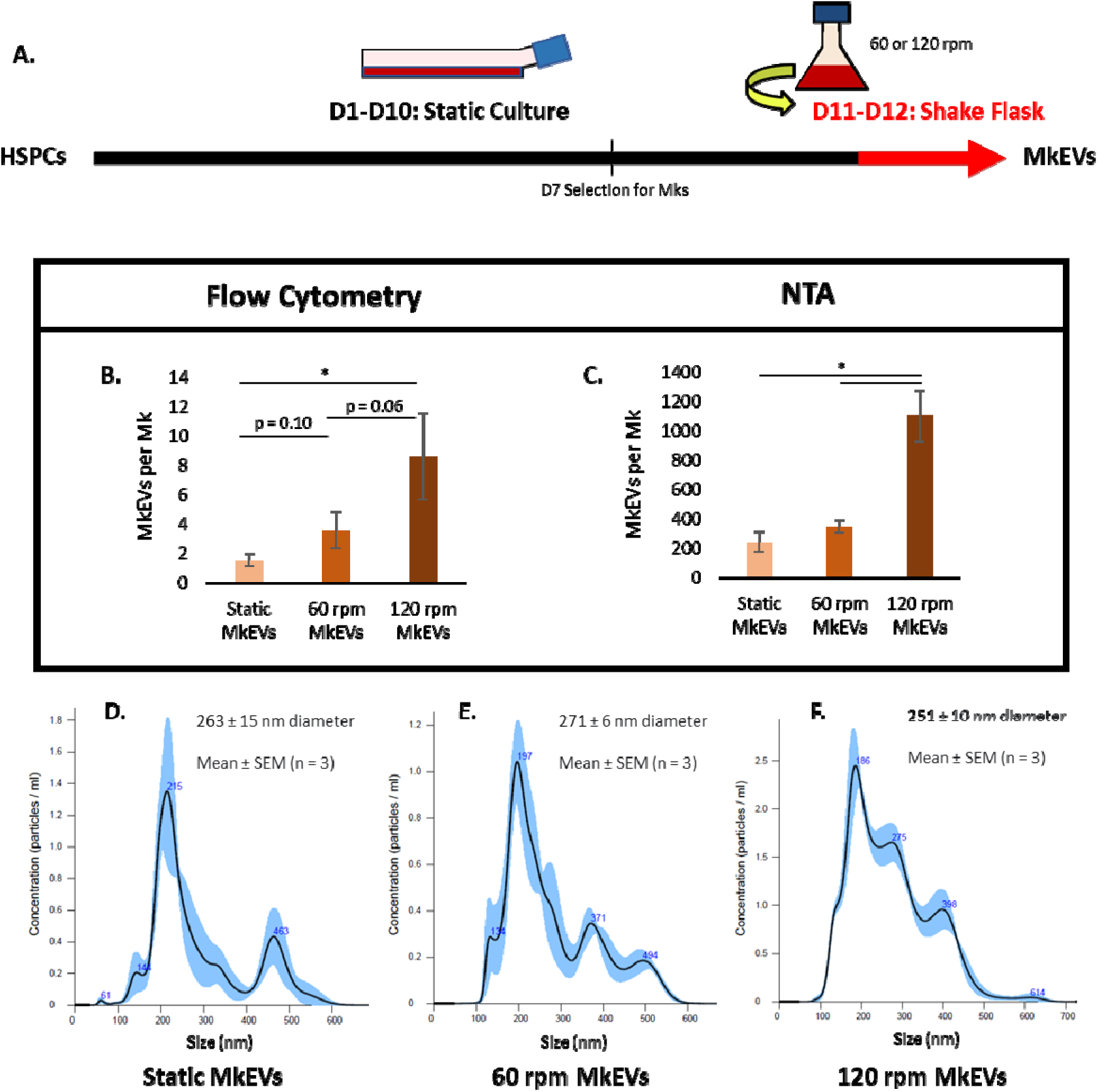
MkEV production rates and size characteristics under complex, long-term biomechanical force. Cells were subjected to rotation in shake flasks at either 60 or 120 rpm during D11 and D12; MkEVs were subsequently collected and isolated. (A) Experimental schematic describing the applied biomechanical force in the context of the overall HSPC/Mk culture process. (B) CD41 MkEV counts, measured via flow cytometry and expressed on a per-Mk basis. (C) MkEV counts, measured via NTA and expressed on a per-Mk basis. (D-F) Sample NTA size distribution profiles for MkEVs from each of the experimental conditions; error bands represent ±1 standard error of the mean (SEM) of three technical replicates. Mean EV diameters and associated SEM values are calculated from three biological replicates. All other error bars indicate SEM of 3-5 biological replicates. Paired Student’s t-tests were performed on all data; * represents p < 0.05.

### 2.3 MkEVs produced under long-term, complex biomechanical forces in shake flasks contain lower levels of two key miRNAs involved in stem cell growth and megakaryocytic differentiation

Total and key individual miRNA levels were measured via Qubit fluorimetry and TaqMan RT-qPCR, respectively. PCR-based quantification of miRNA levels was enabled by use of a spike-in control (cel-miR-39-3p) and subsequent application of the 2^−ΔΔCT^ method,^33^ an increasingly popular technique in the field of EV research.^34,35^ Levels of each miRNA were normalized to both flow cytometry- and NTA-derived MkEV counts. For the shake flask experiments, individual levels of miR-486-5p (Fig. 2A-B) and miR-22-3p (Fig. 2C-D) were significantly higher in MkEVs from static cultures. As before, the flow cytometry-derived results (Fig. 2A, C) and the NTA-derived results (Fig. 2B, D) differed by about 2 orders of magnitude, though trends between samples remained remarkably consistent. Total miRNA levels did not vary significantly between MkEVs, regardless of the MkEV counting method employed (Fig. S1), suggesting that the lower individual miRNA levels observed in the MkEVs produced under high rotational speeds do not reflect a dearth of miRNA content generally. Rather, it affects the sorting and loading of these specific miRNAs into the MkEVs. Sorting and loading of miRNAs in EVs is not a well-understood process, though it is known to involve specific small RNA chaperones.^36–41^

**Figure 2.**
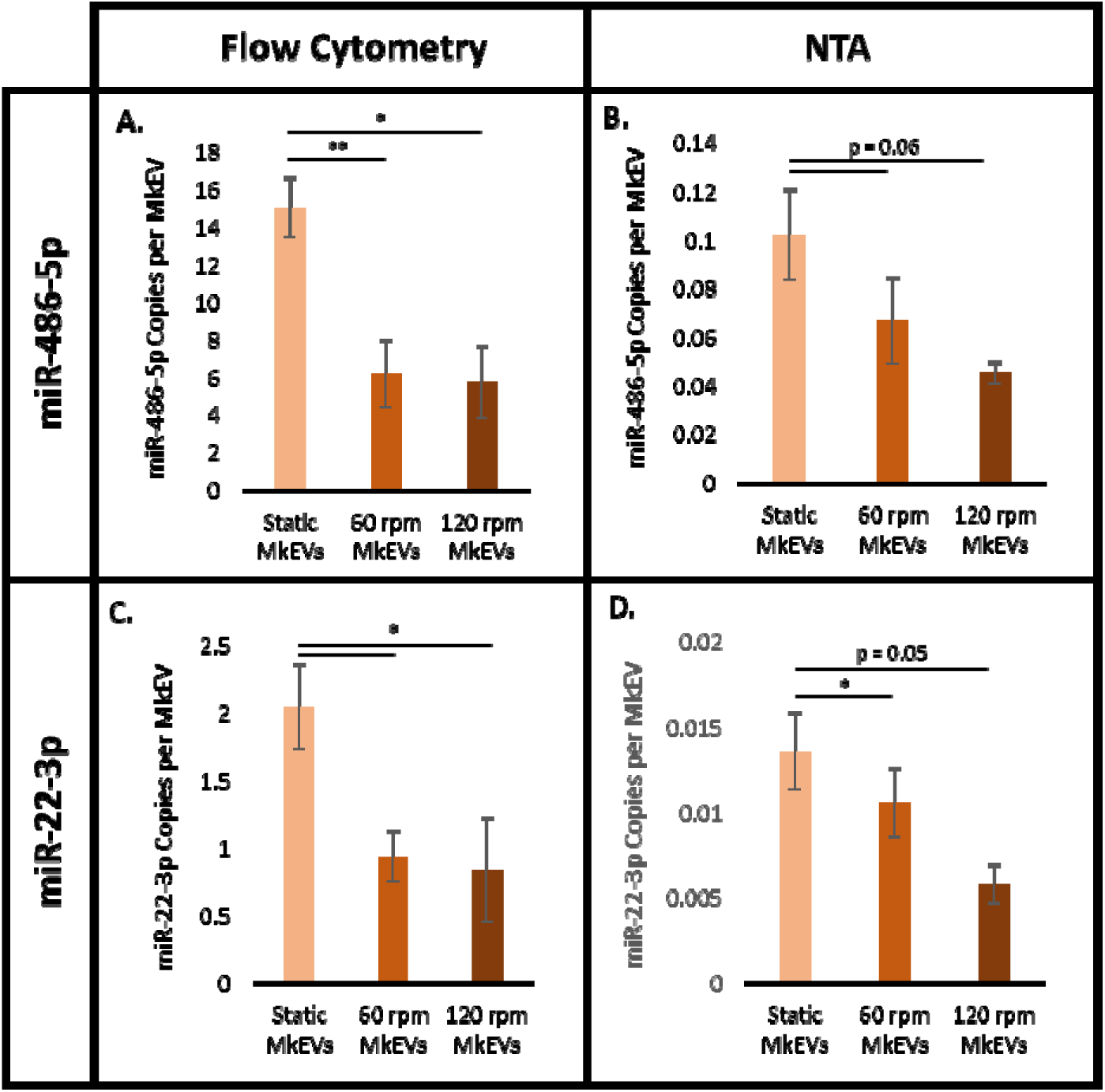
MkEV miRNA content under complex, long-term biomechanical force. (A) Copies of miR-486-5p per MkEV for flow cytometry-based MkEV counts. (B) Copies of miR-486-5p per MkEV for NTA-based MkEV counts. (C) Copies of miR-22-3p per MkEV for flow cytometry-based MkEV counts. (D) Copies of miR-22-3p per MkEV for NTA-based MkEV counts. Error bars indicate SEM of 3 biological replicates. Paired Student’s t-tests were performed on all data; * represents p < 0.05, ** represents p < 0.01.

### 2.4 MkEVs produced under long-term, complex biomechanical forces in shake flasks are less effective in promoting stem cell growth and megakaryocytic differentiation

MkEVs produced in the shake flask experiments were co-cultured with HSPCs for 7 days. In each case, a 20:1 MkEV-to-HSPC ratio (as measured via flow cytometry) was used. TPO treatment was used as a positive control. On day 7 of co-culture, cells were counted (Fig. 3A) and the relative cell fractions expressing CD41 (Fig. 3B) and CD42b (Fig. 3C) were identified. Cell count is expressed as a fold increase relative to untreated HSPC culture. CD41 is a well-known marker for Mks generally (including immature Mks), while CD42b is an early marker for Mk maturation.^13^ The ploidy class distribution (i.e., the fractions of 2N, 4N, and 8N+ cells) for each cell sample was also measured (Fig. 3D). Higher ploidy classes are indicative of late-stage Mk maturation.^1,13^ In general, MkEVs produced under higher levels of shear were relatively less effective, on a per-MkEV basis, at promoting HSPC proliferation (Fig. 3A) and differentiation (Fig. 3B), and this effect was generally dependent on the magnitude of the force (i.e., cell counts and CD41 expression were higher for samples treated with “static” MkEVs than with “60 rpm” MkEVs, and higher for samples treated with “60 rpm” MkEVs than with “120 rpm” MkEVs). While shear-derived MkEVs were likewise less effective in promoting CD42b expression (Fig. 3C), they were similarly effective at promoting high levels of Mk polyploidization (Fig. 3D). Differences in Mk polyploidization may not become apparent until well beyond D7; for this reason, polyploidization was not measured in subsequent co-culture experiments. We suggest that for these shake flask experiments, the reduced efficacy of MkEVs produced under shear could be largely ascribed to differential loading of miRNA cargo, particularly given that the fold changes in individual miRNA cargo levels are comparable to the fold changes in MkEV efficacy (Fig. 2 vs. Fig. 3). Interestingly, the most effective MkEVs promoted Mk maturation better than TPO treatment (as measured by CD42b expression/ploidy class distribution) (Fig. 3C-D), though TPO still maintained an edge as the most effective means of inducing cell proliferation (Fig. 3A), which is consistent with the multifunctional role of TPO in affecting the regulation and proliferation of HSPCs. Cell counts and levels of CD41 and CD42b expression were significantly higher in all MkEV- and TPO-treated cultures than in untreated cultures (not shown), as HSPCs do not spontaneously undergo megakaryopoiesis in any significant numbers. Sample flow cytometry data for CD41/CD42b expression and ploidy distributions are available in Fig. 4.

**Figure 3.**
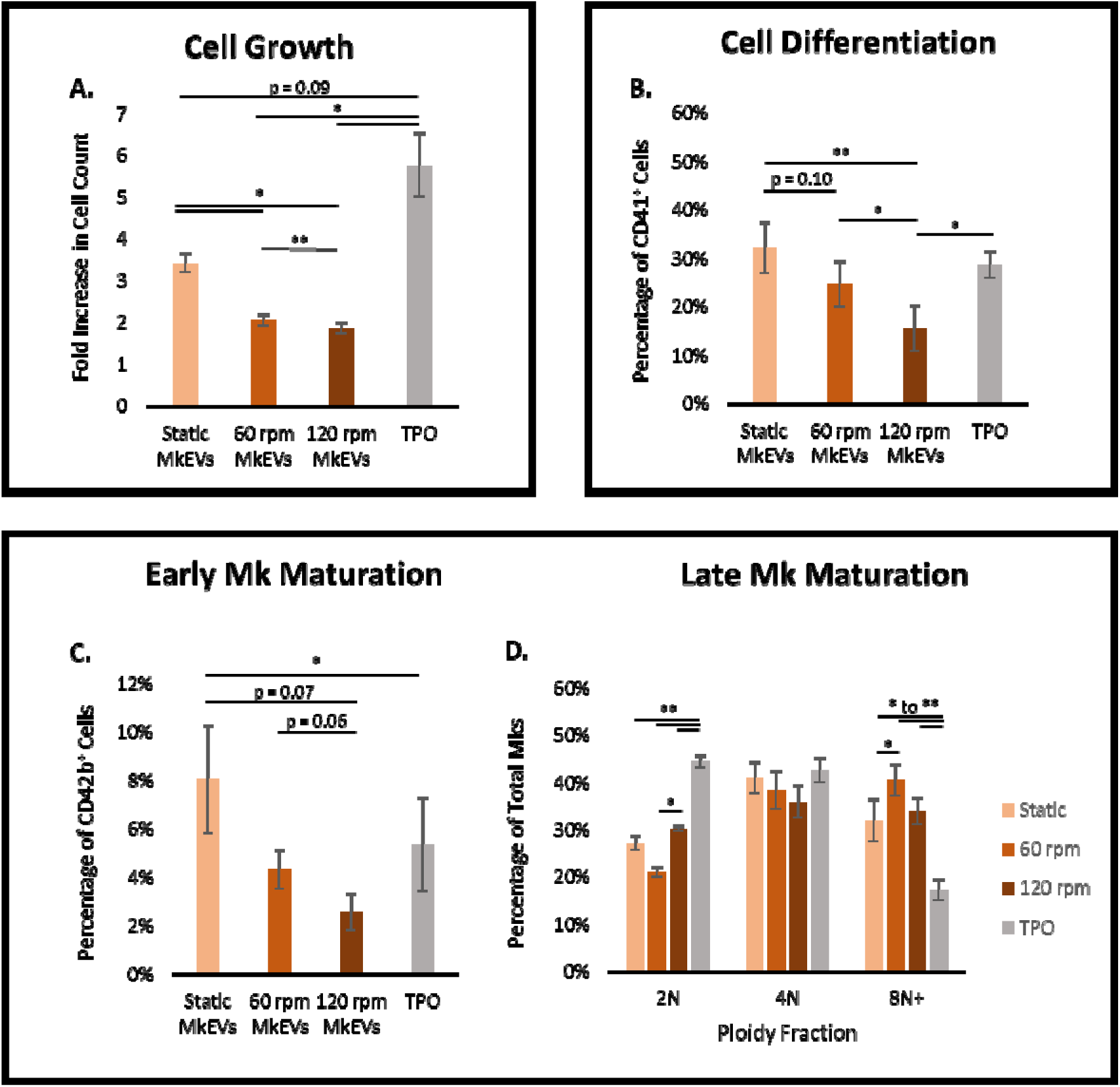
Bioactivity of MkEVs produced under complex, long-term biomechanical force. MkEVs produced under various levels of biomechanical force were co-cultured with HSPCs at a 20:1 ratio for 7 days. (A) Fold change in cell growth (relative to untreated cells) following co-culture with various MkEV samples. (B) The percentage of cells in each co-culture expressing CD41 (an Mk marker). (C) The percentage of cells in each co-culture expressing CD42b (a marker for early Mk maturation). (D) Ploidy fractions for cells in each co-culture; late Mk maturation is associated with higher ploidy numbers. Error bars indicate SEM of 3 biological replicates. Paired Student’s t-tests were performed on all data; * represents p < 0.05, ** represents p < 0.01.

**Figure 4.**
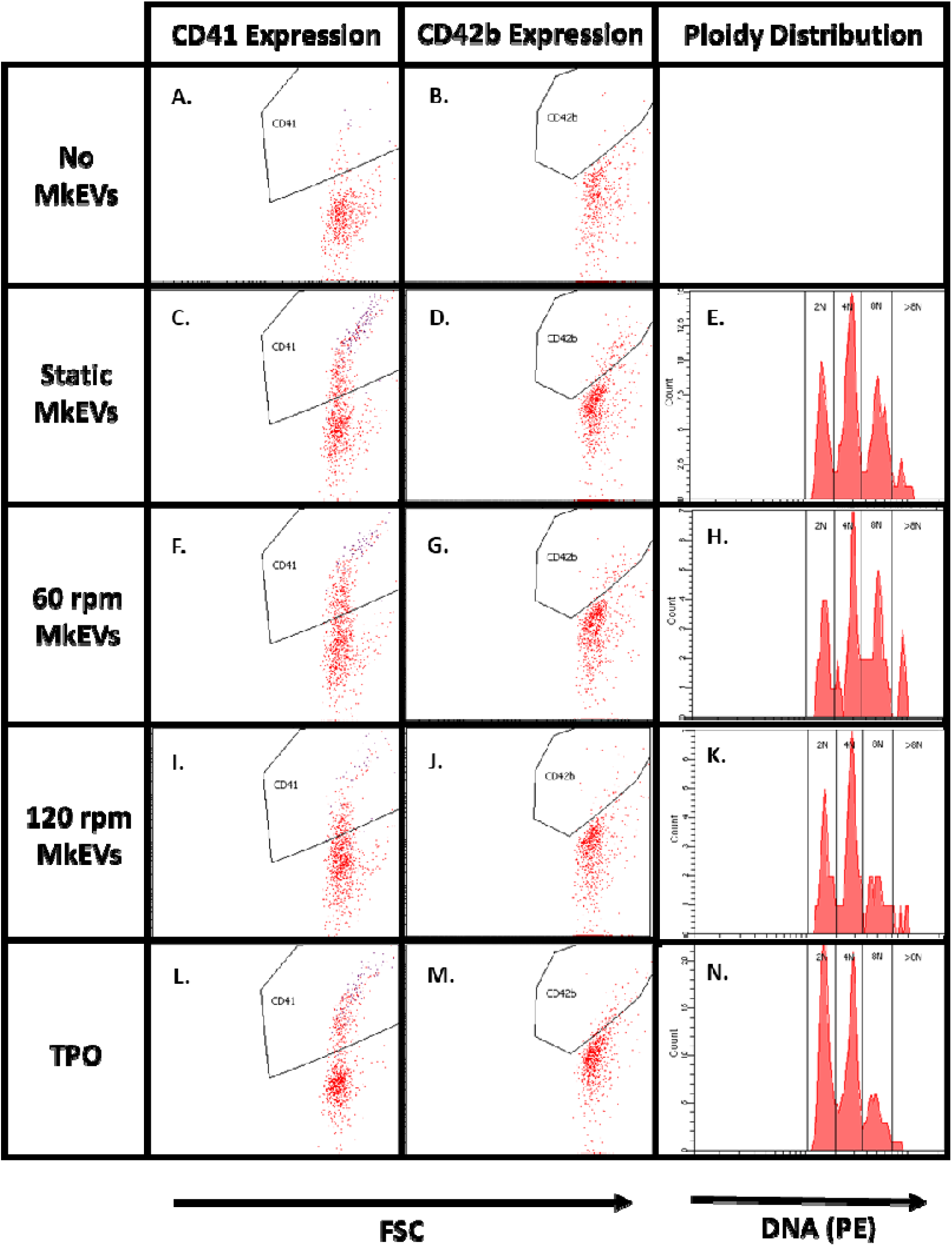
Flow cytometry data for bioactivity of MkEVs produced under complex, long-term biomechanical force. MkEVs produced under various levels of biomechanical force were co-cultured with HSPCs at a 20:1 ratio for 7 days. Sample flow cytometry data are given for cellular CD41 expression, CD42b expression, and Mk ploidy distribution following co-culture with (A-B) no MkEVs or TPO treatment, (C-E) static MkEVs, (F-H) 60 rpm MkEVs, (I-K) 120 rpm MkEVs, and (L-N) TPO treatment.

### 2.5 Brief, high-intensity shear in the syringe pump system increases MkEV production without affecting MkEV size

An experimental schematic for the syringe pump experiments is shown in Fig. 5A. Analysis of MkEVs produced from the syringe pump experiments was similar to the analysis performed on MkEVs from the shake flask experiments. Quantities of MkEVs produced under syringe-induced shear were significantly higher than those produced in static cultures, as evidenced by both flow cytometry (Fig. 5B) and NTA (Fig. 5C) measurements. Here again, the two counting methods differed by roughly 2 orders of magnitude, with MkEV levels under syringe-induced shear comparable to the levels of shake flask-derived MkEVs, despite a 32-fold reduction in shear exposure time (Fig. 5B-C vs. Fig. 1B-C). MkEV size distribution and mean diameter following syringe pump treatment is displayed in Fig. 4D and does not differ significantly from the mean diameter of control MkEVs produced in static conditions (Fig. 5D). Similarly, no significant differences were observed in CD54 or CD11b expression among the MkEV samples (data not shown).

**Figure 5.**
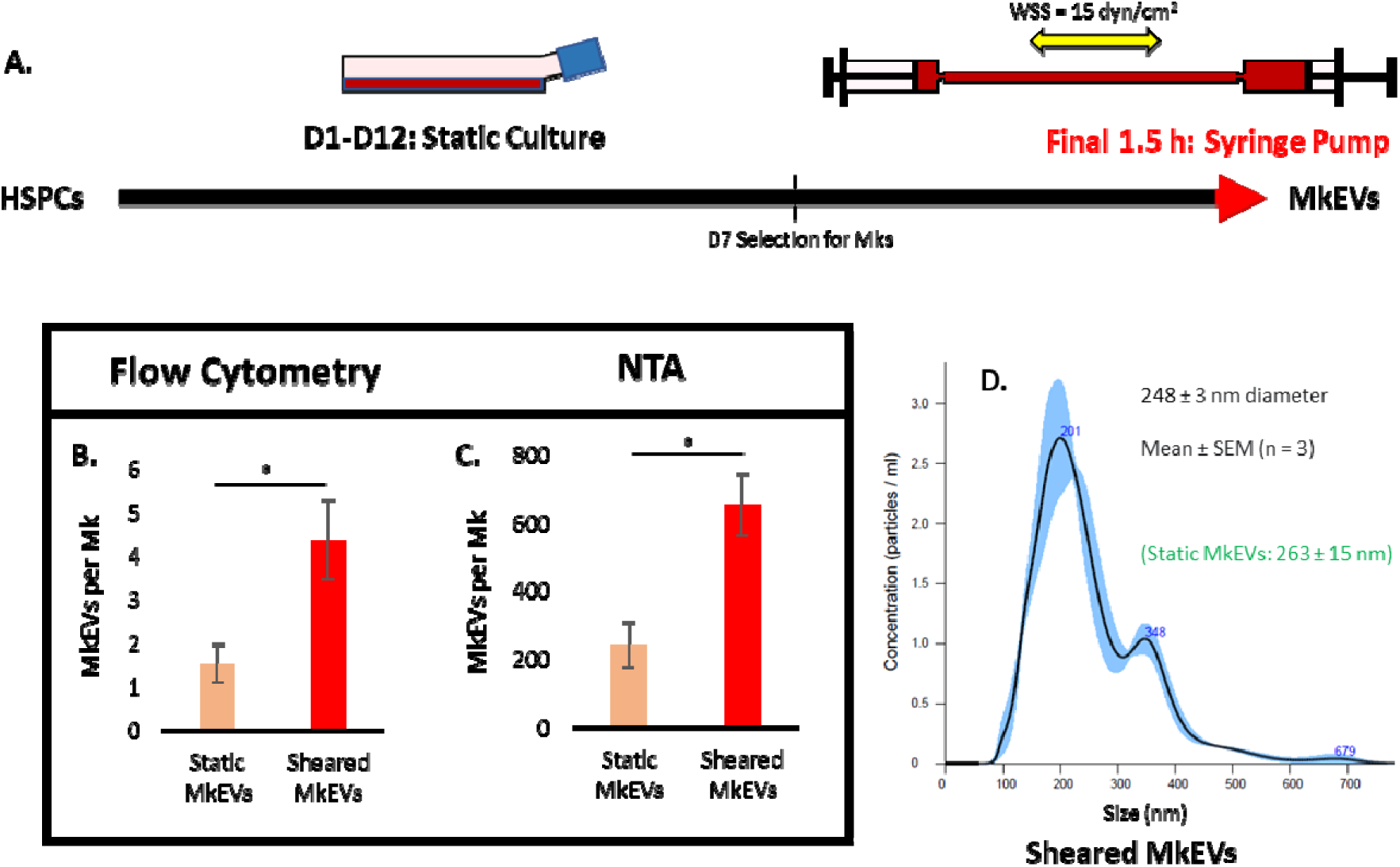
MkEV production rates and size characteristics under brief, defined, and high-intensity biomechanical force. Cells were subjected to 1.5 h of alternating flow in a syringe pump system at the end of D12; MkEVs were subsequently collected and isolated. (A) Experimental schematic describing the applied biomechanical force in the context of the overall HSPC/Mk culture process. (B) CD41 MkEV counts, measured via flow cytometry and expressed on a per-Mk basis. (C) MkEV counts, measured via NTA and expressed on a per-Mk basis. (D) Sample NTA size distribution profile for MkEVs produced in the syringe pump; error bands represent ±1 standard error of the mean (SEM) of three technical replicates. Mean EV diameter and associated SEM value is calculated from three biological replicates. All other error bars indicate SEM of 3-5 biological replicates. Unpaired Student’s t-tests were performed on all data; * represents p < 0.05.

### 2.6 MkEVs produced under brief, high-intensity shear in the syringe pump system are selectively loaded with lower levels of two key miRNAs involved in stem cell growth and megakaryocytic differentiation

Total and key individual (i.e., miR-486-5p and miR-22-3p) miRNA levels in the MkEVs produced under syringe-induced shear were measured as before and compared to the miRNA levels in control MkEVs produced in static conditions. As in the shake flask experiments, individual levels of miR-486-5p (Fig. 6A-B) and miR-22-3p (Fig. 6C-D) were significantly higher in the syringe pump MkEVs than in control MkEVs. Also as before, the flow cytometry-derived results (Fig. 6A, C) and the NTA-derived results (Fig. 6B, D) differed by about 2 orders of magnitude, though trends between samples again remained consistent. Despite varied quantities of individual miRNA cargo, there was no significant difference between total miRNA levels in syringe pump and control MkEVs, regardless of the MkEV counting method employed (Fig. S2).

**Figure 6.**
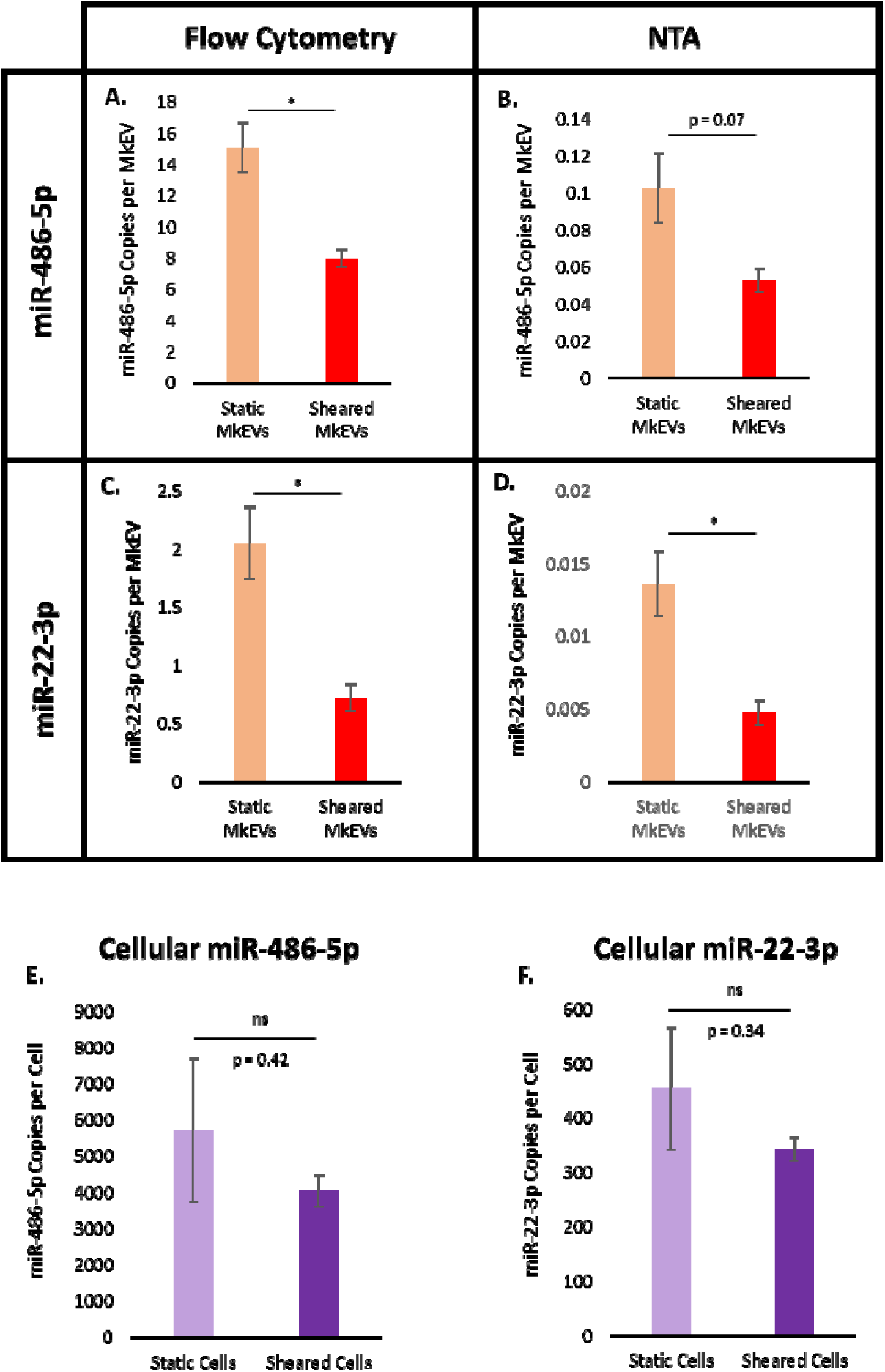
MkEV miRNA content under brief, defined, and high-intensity biomechanical force. (A) Copies of miR-486-5p per MkEV for flow cytometry-based MkEV counts. (B) Copies of miR-486-5p per MkEV for NTA-based MkEV counts. (C) Copies of miR-22-3p per MkEV for flow cytometry-based MkEV counts. (D) Copies of miR-22-3p per MkEV for NTA-based MkEV counts. (E) Copies of miR-486-5p per Mk following syringe pump-induced shear or control treatment. (F) Copies of miR-22-3p per Mk following syringe pump-induced shear or control treatment. Error bars indicate SEM of 3 biological replicates. Unpaired (A-D) or paired (E-F) Student’s t-tests were performed on all data; * represents p < 0.05, ns represents non-significance.

Total and individual miRNA levels were also quantified on a per-cell basis for both the shear-exposed and control Mks; the (individual) miR-486-5p and miR-22-3p levels are shown in Fig. 5E and Fig. 5F, respectively. Notably, there is no significant difference between cellular miRNA levels in either case; combined with the noted differences in the MkEV miRNA levels, these data suggest that miR-486-5p and miR-22-3p cargo is selectively loaded by parent Mks. Indeed, a few quick calculations support this hypothesis. Modeling Mks and MkEVs as spheres and assuming respective diameters of 20 μm and 250 nm (rounded estimates from CellDrop Cell Counter and NTA data), the ratio of Mk volume to MkEV volume is roughly 4 million. Should miRNA cargo be a proportional representation of cellular contents (i.e., “unselectively loaded”), the ratio of cellular miRNA to MkEV miRNA should be similar. However, for static culture, this ratio is roughly 400 or 55,000 (flow cytometry or NTA counting) for miR-486-5p and 200 or 33,000 for miR-22-3p; for syringe pump treatment, this ratio is 500 or 77,000 for miR-486-5p and 500 or 72,000 for miR-22-3p. Thus, miR-486-5p and miR-22-3p loading is highly selective in all experimental conditions, with both individual miRNAs highly upregulated in MkEVs relative to cellular concentrations, even under high-intensity shear. In other words, protein chaperones enrich MkEVs with high concentrations—relative to parent Mks—of the individual miRNAs, though their action appears slightly inhibited by extracellular biomechanical force.

### 2.7 MkEVs produced under brief, high-intensity shear in the syringe pump system possess superior capacity to promote megakaryocytic differentiation and are no less effective than control MkEVs in promoting stem cell growth

MkEVs produced in the syringe pump experiments were co-cultured with HSPCs at a 20:1 ratio for 7 days (as described previously for the shake flask experiments). Once again, on day 7 of co-culture, cells were counted (Fig. 7A) and the relative cell fractions expressing CD41 (Fig. 7B) and CD42b (Fig. 7C) were identified. Sample flow cytometry data for CD41/CD42b expression are shown in Fig. 7D-I. Interestingly, although the syringe pump MkEVs possessed less miR-486-5p and miR-22-3p (Fig. 6), they were no less effective than control MkEVs in promoting cell growth, and were more than twice as effective in promoting megakaryopoiesis (as measured by cellular expression of CD41, Fig. 7B) and Mk maturation (as measured by cellular expression of CD42b, Fig. 7C). The mechanisms underlying this phenomenon are unclear. We suggest that the unique characteristics of the brief, high-intensity shear exerted by the syringe pump imbue the MkEVs with additional miRNA or protein cargo that helps to overcome the reduced efficacy presumably incurred by reductions in the levels of miR-486-5p and miR-22-3p. Indeed, we have previously observed some MkEV-mediated HSPC growth and differentiation even after inhibiting these key miRNAs,^20^ and, as noted in the Introduction, others have identified a possible role for other miRNAs.^21^

**Figure 7.**
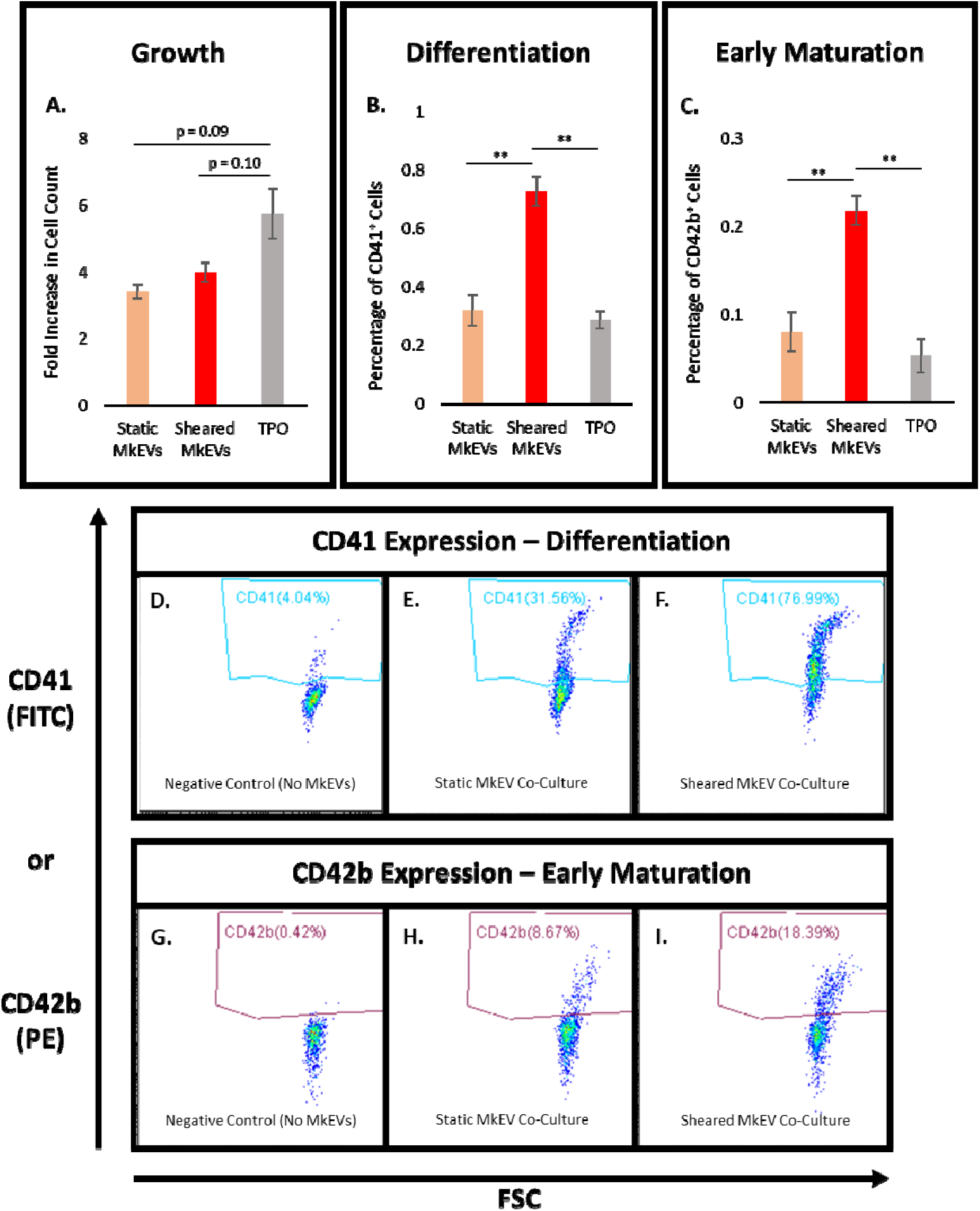
Bioactivity of MkEVs produced under brief, defined, and high-intensity biomechanical force. MkEVs produced in the syringe pump system were co-cultured with HSPCs at a 20:1 ratio for 7 days. (A) Fold change in cell growth (relative to untreated cells) following co-culture with various MkEV samples. (B) The percentage of cells in each co-culture expressing CD41 (an Mk marker). (C) The percentage of cells in each co-culture expressing CD42b (a marker for early Mk maturation). (D-F) Sample flow cytometry data for cellular CD41 expression following (D) co-culture without MkEVs, (E) co-culture with static MkEVs, and (F) co-culture with sheared MkEVs. (G-I) Sample flow cytometry data for cellular CD42b expression following (G) co culture without MkEVs, (H) co-culture with static MkEVs, and (I) co-culture with sheared MkEVs. Error bars indicate SEM of 3 biological replicates. Unpaired Student’s t-tests were performed on all data; * represents p < 0.05, ** represents p < 0.01.

### 2.8 MkEV production increases with Mk culture age, with the MkEVs retaining consistent miRNA levels and bioactivity

Given the impact of biomechanical force on the acceleration of Mk aging and maturation, we hypothesized that Mk age may affect the MkEV quantity and quality in a manner comparable to the shear stress imposed above. MkEVs from D11, D12, and D13 Mk cultures were isolated and counted using both flow cytometry (Fig. 8A) and NTA (Fig. 8B). MkEV levels increased dramatically over this timespan, displaying similar trends regardless of measurement technique, despite NTA counts being higher by the usual 2 orders of magnitude. However, MkEVs from the various days did not display significant differences in mean diameter, CD54/CD11b expression (data not shown), total or individual miRNA levels (Fig. S3), or bioactivity (i.e., ability to induce growth/megakaryopoiesis of HSPCs during co-culture; Fig. S4). Although miR-486-5p levels per MkEV did appear to dip slightly from D11 to D13 when MkEVs were quantified using flow cytometry, this finding was not supported by calculations employing NTA-derived MkEV counts (Fig. S3C-D).

**Figure 8.**
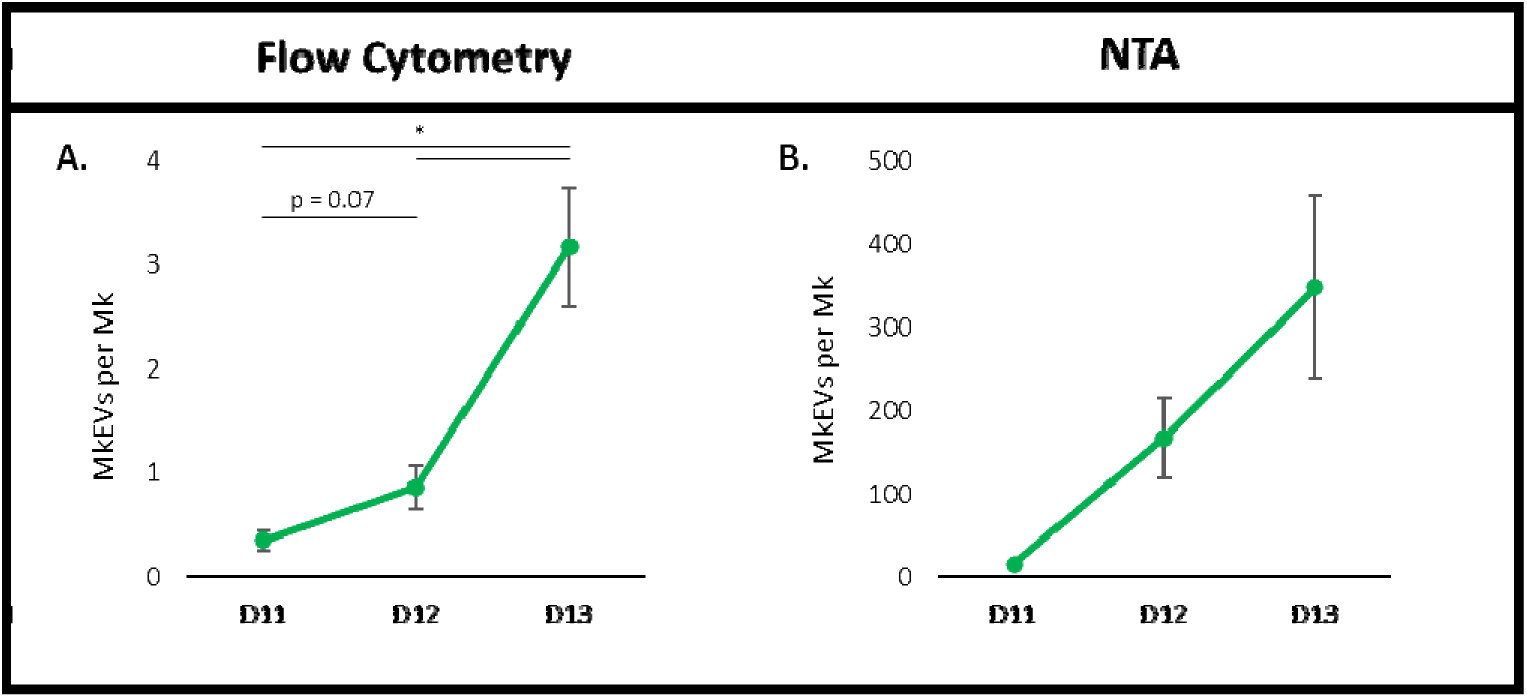
Kinetics of MkEV production by mature Mks. MkEVs from D11-D13 Mks were quantified via (A) flow cytometry and (B) NTA; counts are expressed on a per-Mk basis. Error bars indicate SEM of 3-4 biological replicates, except in the case of NTA-derived D11 MkEV counts (1 biological replicate). Paired Student’s t-tests were performed on (A) only; * represents p < 0.05.

### 2.9 Delayed HSPC differentiation results in Mks with reduced capacity for MkEV production

Given the impact of Mk age on MkEV production, we hypothesized that delayed differentiation of HSPCs (into Mks) may also impact Mk productivity and, therefore, MkEV production. As described previously, CD34^+^ cells (undifferentiated HSPCs) were continually re-cultured, such that Mks and their MkEVs could be lumped into three categories: those arising from HSPCs that underwent megakaryopoiesis between D1 and D7 (“D1-D7 differentiation”), those arising from HSPCs that underwent megakaryopoiesis between D8 and D14 (“D8-D14 differentiation”), and those arising from HSPCs that underwent megakaryopoiesis between D15 and D21 (“D15-D21 differentiation”). An experimental schematic is provided in Fig. 9A-B. Mk production for each category was quantified on a per-HSPC basis and plotted in Fig. 9C. Similarly, MkEV production for each category was quantified (using flow cytometry) on a per-Mk basis and plotted in Fig. 9D. Taken together, the data suggest that as time passes and HSPCs age and replicate, they are progressively less likely to undergo megakaryopoiesis, and the Mks they do produce are increasingly ineffective at producing MkEVs. We also plotted total Mk and MkEV production per initial (thawed) HSPC as a function of time (Fig. 9E-F). Though numeric values were variable due to one highly productive outlier, trends were consistent: both Mk and MkEV production increased substantially following the first “recycle” of CD41^−^/CD61^−^ cells, but not following the second “recycle.” From a biomanufacturing perspective, this suggests that attempts to continue deriving Mks/MkEVs from a given HSPC culture after ~19 days will prove highly inefficient.

**Figure 9.**
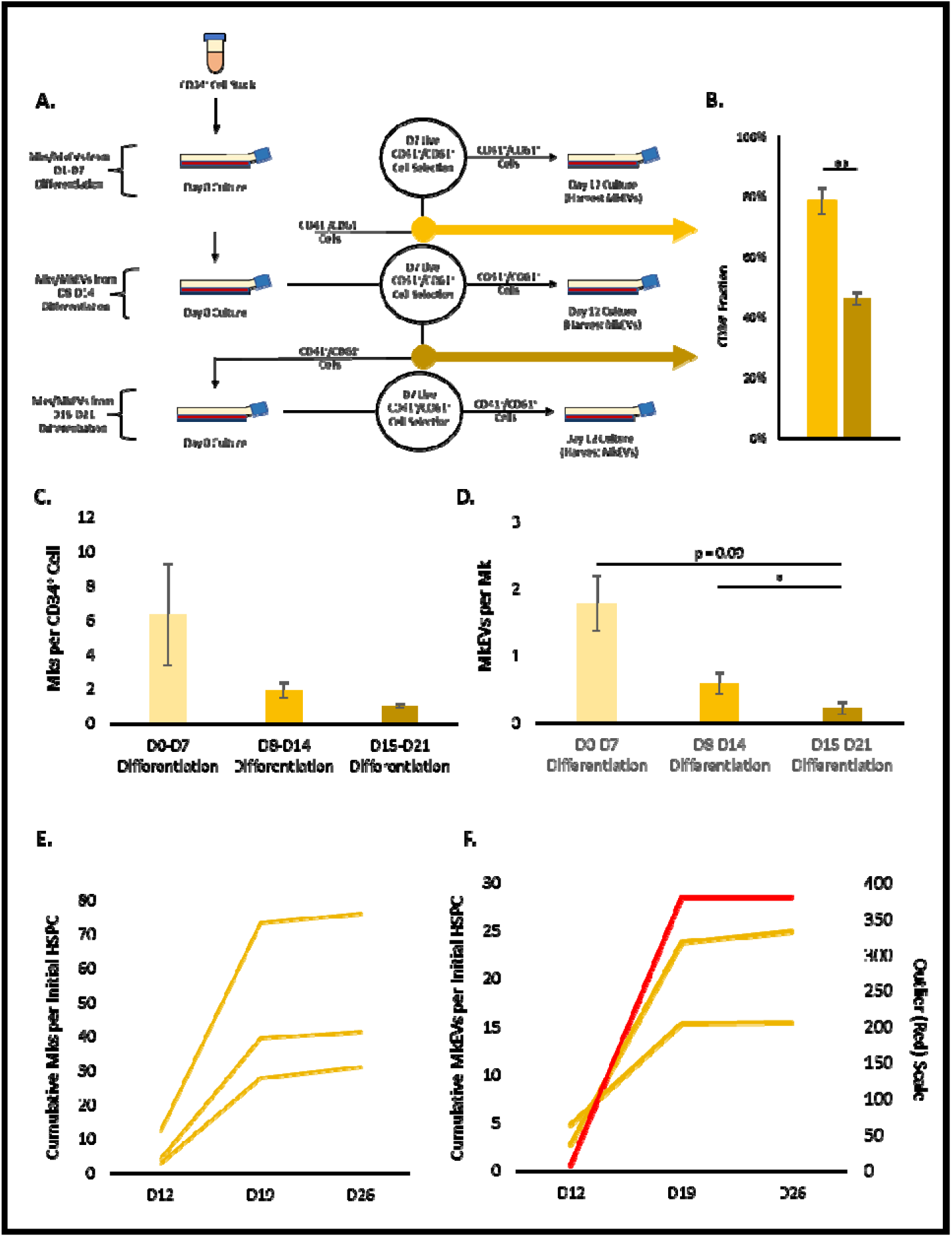
MkEV production as a function of HSPC differentiation time. Mks were cultured using standard protocols. Undifferentiated (CD41 /CD61) cells from each Mk selection were re-cultured. (A) An experimental schematic describing the culture process and the re-culture of undifferentiated cells following each Mk enrichment process. (B) The percentage of cells expressing CD34 in each sample of undifferentiated cells; the colored arrows correspond to the bars on the graph (i.e., the lighter color represents CD34 expression in the first generation of undifferentiated cells, while the darker color represents CD34 expression in the second generation of undifferentiated cells. (C) The average number of Mks produced per CD34 cell by the original culture, the first generation of undifferentiated cells, and the second generation of undifferentiated cells. (D) The average number of MkEVs produced by Mks from the original culture, the first generation of undifferentiated cells, and the second generation of undifferentiated cells. (E) Cumulative Mks produced per initial HSPC after 26 days. Three biological replicates are represented as three individual trendlines. (F) Cumulative MkEVs produced per initial HSPC after 26 days. Three biological replicates are represented as three individual trendlines; one outlier (shown in red) uses a different y-axis. Error bars indicate SEM of 3 biological replicates. Paired Student’s t-tests were performed on A-D; * represents p < 0.05.

## 3 DISCUSSION

As was once the case with cellular protein glycosylation, EV quality is increasingly recognized as culture- and process-dependent, with numerous—and often overlooked—variables impacting product efficacy. Perhaps the most notable of these variables is biomechanical force, which is ubiquitous in biomanufacturing and therefore vital for the clinical implementation of EV technology. Other variables of note include culture age, media composition and pH, and oxygen tension. Beyond simply expanding the rate of EV production, biomanufacturing processes must ensure that EV quality is standardized. Surface protein expression, miRNA and protein cargo levels, and morphological characteristics including size must all be consistently maintained if EVs are to be successfully harnessed as therapeutics.

This study highlights numerous novel impacts of Mk culture conditions on MkEV quantity and quality. Specifically, relative to static controls, Mks exposed to various types, magnitudes, and durations of biomechanical force produced higher numbers of MkEVs containing lower levels of miR-486-5p and miR-22-3p miRNA cargo. Possibly as a result, some of these MkEVs—those derived under long-term, mild shaking—were relatively less efficacious (in terms of their ability to spur growth and megakaryopoiesis of HSPCs). However, the efficacy of MkEVs derived under the brief, high-intensity shear of the syringe pump matched or exceeded that of control MkEVs. Therefore, where therapeutic applications (e.g., alleviating thrombocytopenia) are desirable, brief, high-intensity shear offers respite from the tradeoffs inherent in long-term, mild shaking, producing MkEVs that are both numerous and efficacious, despite a relative depletion of key miRNA cargo. This paper also notes significant increases in MkEV production from D11 to D13, as parent Mks age and mature. However, in contrast with biomechanical force, culture age did not affect MkEV miRNA levels or efficacy. At the same time, Mks arising from older HSPCs that exhibited delayed differentiation were less productive and yielded fewer MkEVs.

Changes in miRNA cargo levels and subsequent EV phenotypes in response to variations in biomechanical force are well-documented for a wide variety of cell types, including endothelial cells,^42,43^ fibroblasts,^44^ muscle cells,^45,46^ bone cells,^47,48^ Schwann cells,^49^ bronchial epithelial cells,^50,51^ and mesenchymal stem cells (MSCs).^52^ However, this paper represents the first such documentation of shear-induced miRNA cargo variations in MkEVs.

We distinguish between two different hypotheses for the relative dearth of miR-486-5p and miR-22-3p in MkEVs produced under shear. In the first, miRNA loading machinery is unaffected by shear, but simply cannot keep pace with the increased rate of plasma membrane shedding triggered by the extracellular force. In the second, shear stress triggers signaling cascades that fundamentally alter the biology of the Mks, and, subsequently, their miRNA loading machinery. While the second hypothesis is a feasible explanation for the impact of long-term shear, we question whether short-term shear (e.g., the 1.5 h timeframe employed in the syringe pump experiments) can meaningfully change the EV loading machinery. Moreover, unlike more compact cells, mature Mks are uniquely susceptible to biomechanical forces as a result of their wispy proplatelet extensions,^4^ suggesting the first hypothesis may partly explain MkEV production in particular, as MkEVs are derived directly from the plasma membrane. Under static conditions, however, MkEV production occurs independently of proplatelet formation and is instead reliant on actin depolymerization.^53^ We have already demonstrated an extreme manifestation of the first hypothesis, wherein Mks are extruded with such force as to rip apart the plasma membrane, creating a multitude of membrane fragments which can spontaneously reassemble to form novel “empty” EVs with a normal landscape of surface proteins.^54^ In the end, the most likely explanation for shear-induced variation in MkEV cargo is one that includes both hypotheses and encompasses numerous mechanisms, all of which vary with shear duration and magnitude. Indeed, in cultured endothelial cells, shear level has been found to mediate not only MP production rate, but also the mechanism by which production occurs;^55^ a similar phenomenon may occur where MkEV cargo loading is concerned.

The impacts of culture age and delayed HSPC differentiation on MkEV quantity and quality have been hereto unexamined, and the general relationship between cell age and EV quality has been studied only sparingly in other cell types. MSC-derived EVs display notable changes in cargo and reductions in function as their parent cells become senescent.^56^ Relative to EVs from fresh (day 1) platelets, EVs from platelets stored for 5 days are more abundant, enriched in long-chain ceramide, and depleted of sphingosine-1-phosphate, inducing lung injury in vivo as a result.^57^ Among Chinese hamster ovary (CHO) cells, too, EV production rate varies with culture age,^58^ and we have recently discovered age-dependent variation in CHO MP cargo, as well (manuscript in preparation). The similarities in MkEV yields between “older” cultures and high-shear cultures are substantial, despite differences in MkEV miRNA cargo and efficacy. Significantly, as noted above, we have previously identified a role for shear stress in promoting early Mk maturation.^13,28^ The noted similarities between “older” and high-shear cultures may therefore arise partly from this precise mechanism; that is, the primary effect of shear may be to “age” Mks, prompting them to produce higher numbers of MkEVs and mimicking the observed boost in “late-stage” MkEV production by older cultures.

Controversy exists as to the “true” miRNA levels in EVs. Some prior research has identified as little as 1 copy of abundant miRNA species for every 10 to 100 EVs.^59,60^ However, another study has noted much higher miRNA concentrations: 12 to 63 copies of abundant miRNA species in just a single EV.^61^ One paper of particular note analyzed only large EV fractions (i.e., MPs sorted using flow cytometry) and identified 25 to 35 copies of abundant miRNA species in each particle.^34^ A rough approximation of volume in a 100 nm EV also suggests capacity for up to 1,000 miRNA-protein complexes.^34^ While our flow cytometry-based MkEV counts suggest miRNA levels similar to these higher values, our NTA-based MkEV counts reflect much lower miRNA levels. The reality is probably somewhere in the middle. While traditional flow cytometry (used in this study) is ineffective at detecting small (i.e., <200 nm) particles, NTA counts are inflated by non-EV particles such as protein aggregates and lipoproteins.^32^ Larger EVs are also capable of carrying larger miRNA loads, and our ultracentrifugation protocol enriches for large EVs (i.e., MPs) while eliminating a significant portion of small EVs (i.e., Exos).

Future investigations should identify the cargo responsible for the enhanced efficacy of syringe pump-derived MkEVs, a project which will likely require RNA sequencing similar to that previously conducted by our group.^20^ Researchers should also aim to elucidate the particular mechanisms by which miRNA is loaded into MkEVs. While the general nature of EV cargo loading is receiving significant new attention and has been well-reviewed elsewhere,^36–41^ the landscape of mechanisms is diverse, and nothing is known regarding Mks specifically. A thorough understanding of EV loading machinery in Mks would aid in explaining the phenomena characterized in this paper. An investigation into the effects of individual forces (e.g., shear, tension, and/or compression) on MkEV production may also prove fruitful. Although these forces are almost never isolated in vivo or among the complex flows of sparged or stirred bioreactors, they nonetheless often affect EV production in different ways, even among similar cell types. For instance, with increasing shear stress, human umbilical vein endothelial cells (HUVECs) produced fewer MPs,^55^ while similar human pulmonary artery endothelial cells (HPAECs) subjected to cyclic stretching produced far greater numbers of MPs than their unstretched counterparts.^62^ A similar phenomenon may be responsible for the enhanced efficacy of syringe pump-derived MkEVs. Finally, as the literature surrounding MkEVs continues to emerge, future studies should aim to determine the impact of culture parameters such as biomechanical force or Mk age on any newly-discovered MkEV functions or cargo. For instance, one recent study suggests that MkEVs influence the progression of arthritis via the delivery and release of inflammatory cytokine IL-1.^63^ Understanding the influence of physiological stressors on IL-1 loading (into MkEVs) may therefore accelerate the development of new arthritis treatments.

## 4 CONCLUSION

In sum, we present a novel examination of the ways in which key culture parameters— biomechanical force, Mk age, and delayed HSPC differentiation—impact MkEV quantity and quality, highlighting a path toward the continued correlation of MkEV characteristics with key scalable production parameters and thereby providing a framework that links laboratory know-how with clinical and industrial possibilities. We hope that our methodology—which links culture conditions with EV production rate, genotype, and phenotype—will become increasingly utilized as necessary in the development of biomanufacturing strategies, for although EVs produced under different conditions may appear similar, their underlying properties are often highly distinct.

## 5 MATERIALS AND METHODS

### 5.1 Chemicals, reagents, and antibodies

Iscove’s Modified Dulbecco’s Medium (IMDM) was purchased from Gibco (Waltham, MA, USA). BIT 9500 Serum Substitute was purchased from STEMCELL Technologies (Vancouver, BC, Canada). Recombinant human TPO, stem cell factor (SCF), and interleukins 3 (IL-3), 6 (IL-6), 9 (IL-9), and 11 (IL-11) were purchased from PeproTech (Cranbury, NJ, USA). CD61 MicroBeads and other MACS cell separation equipment were purchased from Miltenyi Biotec (Bergisch Gladbach, Germany). Human LDL (hLDL) was purchased from Sigma-Aldrich (St. Louis, MO, USA). Fluorescein isothiocyanate (FITC)-conjugated anti-CD41, phycoerythrin (PE)-conjugated anti-CD42b, PE-conjugated anti-CD54, and allophycocyanin (APC)-conjugated anti-CD11b antibodies were purchased from BD Biosciences (Franklin Lakes, NJ, USA). Unless otherwise noted, all other chemicals were obtained from Thermo Fisher Scientific (Waltham, MA, USA) or Sigma-Aldrich (St. Louis, MO, USA).

### 5.2 Megakaryocyte culture

Frozen G-CSF (granulocyte colony-stimulating factor)-mobilized human peripheral blood CD34^+^ cells (i.e., HSPCs) were obtained from the Fred Hutchinson Cancer Research Center (Seattle, WA) and stored in liquid N_2_. HSPC/Mk culture proceeded according to previous protocols^11,13^ developed from optimization experiments by our group.^2,3^ Briefly, cells were suspended in IMDM media supplemented with BIT 9500, TPO, SCF, IL-3, IL-6, IL-11, and hLDL and cultured in 5% O_2_. After 5 days, cells were suspended in a new, similar media cocktail that replaced IL-6 with IL-9 and were cultured thereafter in 20% O_2_. After 7 days, dead cells were removed from the culture via a Miltenyi Biotec Dead Cell Removal Kit and MiniMACs Separator with accompanying columns (Miltenyi Biotec, Bergisch Gladbach, Germany). CD61^+^ Mks were then enriched from the live cells using CD61 MicroBeads and a MidiMACs Separator with accompanying columns (Miltenyi Biotec, Bergisch Gladbach, Germany). The resulting sample of live CD61^+^ Mks was resuspended in IMDM media supplemented with BIT 9500, TPO, SCF, hLDL, and nicotinamide. Except in studies of Mk age, cultures were ended and MkEVs were isolated (as described below) after 12 total days. Where necessary, cell counts were obtained using a CellDrop Cell Counter (DeNovix, Wilmington, DE, USA).

Certain alterations to the described culture procedures were implemented as appropriate. For the shake flask experiments, Mks were transferred (without any media change) from T-flasks to shake flasks after 10 total days and agitated on shaker plates as described (i.e., at either 60 or 120 rpm). For the experiments examining Mk age, MkEVs were collected after either 11, 12, or 13 total days. For the experiments involving delayed HSPC differentiation, CD41^−^/CD61^−^ cell samples collected on D7 were resuspended in D0 media and treated as a D0 culture; this process was then repeated one additional time.

### 5.3 Syringe pump production

For the syringe pump experiments, day 12 Mks in standard culture media were loaded into 50 mL syringes (BD Biosciences, Franklin Lakes, NJ, USA) connected by 250 mm of sterilized, 1.58 mm ID Fisherbrand traceable silicone pump tubing (Thermo Scientific, Waltham, MA, USA). The connected syringes were secured in a Dual NE-4000 Two Channel Programmable Syringe Pump (New Era Pump Systems, Farmingdale, NY, USA) and alternately discharged into one another at a flow rate of 4,448 mL/h for 1.5 h. This flow rate corresponded to a wall shear stress of 15 dyn/cm^2^ in the connective tubing. After 1.5 h, MkEVs were isolated from the media according to the procedure below. Cell viability was measured via trypan blue staining and a CellDrop Cell Counter (DeNovix, Wilmington, DE, USA).

### 5.4 Isolation of MkEVs

MkEVs were isolated as described previously.^11,13^ Briefly, apoptotic bodies and platelet-like particles were pelleted by centrifugation at 2,000 × g for 10 min. and discarded. The resulting supernatant was subjected to ultracentrifugation at 25,000 × g for 30 min. at 4°C using an Optima Max Ultracentrifuge and a TLA-55 rotor (Beckman Coulter, Brea, CA, USA). The pellet from this supernatant (i.e., the MkEVs) was resuspended in PBS or pure IMDM media and frozen at −80°C until further use.

### 5.5 Quantification and flow cytometry of MkEVs

MkEVs were analyzed using flow cytometry as described previously.^11,13^ Briefly, MkEVs were stained with FITC-conjugated CD41 antibody prior to counting via flow cytometry. Samples were analyzed with either a BD FACSAriaII flow cytometer using FACSDiva software (BD Biosciences, Franklin Lakes, NJ, USA) or a CytoFLEX flow cytometer using CytExpert software (Beckman Coulter, Brea, CA, USA). For the BD FACSAriaII, a known (10 uL) quantity of ~5.0 um, 10^6^/mL AccuCount Fluorescent Beads (Spherotech, Lake Forest, IL, USA) was added to each sample for MkEV quantification. MkEVs were characterized as CD41^+^ particles of between 0.2 um and 1 um in diameter, with size limits determined using fluorescent calibration beads from Spherotech (Lake Forest, IL, USA). For surface protein examination, MkEVs were simultaneously stained with additional PE- or APC-conjugated CD54 or CD11b antibodies, respectively (described above).

### 5.6 Size distribution of MkEVs

MkEVs were diluted by between 25- and 35-fold in filtered PBS and characterized using a NanoSight NS300 instrument (Malvern Panalytical, Malvern, United Kingdom) as described previously.^18^ For each biological replicate, measurements were taken in triplicate and averaged.

### 5.7 HSPC-MkEV co-cultures

Co-culture procedures closely mimicked those described previously.^13^ MkEVs and HSPCs suspended in pure IMDM were combined in microcentrifuge tubes at predetermined ratios and subsequently incubated under 20% O_2_ for 1 h. MkEV-to-HSPC ratios of 20:1 were used. MkEV and HSPC counts were obtained via flow cytometry/NTA and a CellDrop Cell Counter, respectively. Total co-culture volumes were kept constant (<50 uL) for each experiment by adding pure IMDM as appropriate. Following the first hour of co-culture, samples were transferred to well plates and diluted to densities of 100,000 cells/mL using a mixture of IMDM supplemented with 5% BIT 9500 and 50 ng/mL SCF. The use of lower volumes for initial (1 h) co-culture allows for greater contact between cells and MkEVs.^13^ Cultures were analyzed after 7 days.

### 5.8 Flow cytometry of Mks

Mks were analyzed using flow cytometry as described previously.^11,13^ Briefly, Mks were stained with FITC-conjugated CD41 antibody and PE-conjugated CD42b antibody. Samples were analyzed using the flow cytometers and associated software/methods described above.

### 5.9 Ploidy analysis of Mks

Ploidy analysis was performed as described previously.^64^ Briefly, Mks were stained with FITC-conjugated CD41 antibody, fixed with 0.5% paraformaldehyde in PBS, and permeabilized with 70% methanol. RNA was eliminated via RNase treatment and DNA was stained with 50 ug/mL propidium iodide in PBS. Samples were analyzed using the flow cytometers and associated software/methods described above. Ploidy fractions were calculated by identifying the number of CD41^+^ cells with either 2N, 4N, or ≥8N DNA content.

### 5.10 Isolation of miRNA

Isolation of miRNA occurred as described previously.^20^ Briefly, HSPCs, Mks, or MkEVs were pelleted via centrifugation or ultracentrifugation (as described above). RNA extraction was performed using the miRNeasy Micro Kit (QIAGEN, Hilden, Germany) according the manufacturer’s instructions. For RT-PCR analysis, 0.5 pmol synthetic cel-miR-39-3p (Thermo Scientific, Waltham, MA, USA) was added to the samples during cell/MP lysis as a spike-in control. Extracted RNA was flash frozen in liquid N_2_ and stored at −80°C.

### 5.11 Quantification of total miRNA

Isolated miRNA samples were processed using a Qubit miRNA Assay Kit (Thermo Scientific, Waltham, MA, USA), and subsequently analyzed using a Qubit 3.0 Fluorometer (Thermo Scientific, Waltham, MA, USA) according to the manufacturer’s instructions.

### 5.12 Quantitative RT-PCR

Quantification and RT-PCR of key miRNAs occurred as described previously.^20^ Reverse transcription of extracted miRNA was performed using the ThermoFisher TaqMan MicroRNA Reverse Transcription Kit and the appropriate primers and probes (Thermo Scientific, Waltham, MA, USA) according to the manufacturer’s instructions. Subsequent PCR was performed using TaqMan Universal PCR Master Mix II and the appropriate TaqMan Small RNA Assays (Thermo Scientific, Waltham, MA, USA), again according to the manufacturer’s instructions. PCR for each biological replicate was performed in triplicate. The CFX96 Optical Reaction Module (Bio-Rad, Hercules, CA, USA) was used to control the PCR reactions. Calculation of miRNA levels employed the 2^−ΔΔCT^ method.^33^

### 5.13 Statistical analysis

Data are presented as mean(s) ± one standard error of the mean(s), with each mean informed by ≥3 biological replicates. Student’s t-test (paired or unpaired, as appropriate) was performed for all data, with statistical significance indicated as follows: * for p < 0.05, ** for p < 0.01, and ns for notable cases of non-significance.

### 5.14 Institutional Review Board statement

This work adhered to the Helsinki Declaration of 1975 and was approved by the Institutional Review Board at the University of Delaware.

## CRediT authorship contribution statement

Will Thompson: conceptualization, investigation, formal analysis, writing - original draft, writing - review & editing. Eleftherios Terry Papoutsakis: conceptualization, funding acquisition, writing - review & editing, supervision.

## Data availability

The data supporting the findings of this study are available from the corresponding author upon reasonable request.

## Abbreviations

Akt: protein kinase B
APC: allophycocyanin
BIT: bovine serum albumin, insulin, and transferrin
CD: cluster of differentiation
CHO: Chinese hamster ovary
CXCR4: C-X-C chemokine receptor type 4
D#: day #
EV: extracellular vesicle
Exos: exosome
FITC: fluorescein isothiocyanate
G-CSF: granulocyte colony-stimulating factor
hLDL: human low-density lipoprotein
HPAEC: human umbilical vein endothelial cell
HSPC: hematopoietic stem and progenitor cell
HUVEC: human pulmonary artery endothelial cell
IL: interleukin
IMDM: Iscove’s Modified Dulbecco’s Medium
JNK: c-Jun N-terminal kinase
MACS: magnetic-activated cell sorting
miRNA: microRNA
Mk: megakaryocyte
MkEV: megakaryocyte-derived extracellular vesicle
MkExo: megakaryocyte-derived exosome
MkMP: megakaryocyte-derived microparticle
MP: microparticle
MSC: mesenchymal stem cell
mTOR: mammalian/mechanistic target of rapamycin
MVB: multivesicular body
ns: not significant
NTA: nanoparticle tracking analysis
PBS: phosphate-buffered saline
PCR: polymerase chain reaction
PE: phycoerythrin
PI3K: phosphoinositide 3-kinase
qPCR: quantitative polymerase chain reaction
rH: relative humidity
rpm: revolutions per minute
RT: reverse transcription
SCF: stem cell factor
TPO: thrombopoietin
WSS: wall shear stress

## REFERENCES

1. Machlus KR, Italiano JE, Jr. The incredible journey: From megakaryocyte development to platelet formation. J Cell Biol. 2013;201(6):785–796. doi:10.1083/jcb.201304054

2. Panuganti S, Papoutsakis ET, Miller WM. Bone marrow niche-inspired, multiphase expansion of megakaryocytic progenitors with high polyploidization potential. Cytotherapy. 2010;12(6):767–782. doi:10.3109/14653241003786148

3. Panuganti S, Schlinker AC, Lindholm PF, Papoutsakis ET, Miller WM. Three-stage ex vivo expansion of high-ploidy megakaryocytic cells: toward large-scale platelet production. Tissue Eng Part A. 2013;19(7-8):998–1014. doi:10.1089/ten.TEA.2011.0111

4. Junt T, Schulze H, Chen Z, et al. Dynamic visualization of thrombopoiesis within bone marrow. Science. 2007;317(5845):1767–1770. doi:10.1126/science.1146304

5. Lefrancais E, Ortiz-Munoz G, Caudrillier A, et al. The lung is a site of platelet biogenesis and a reservoir for haematopoietic progenitors. Nature. 2017;544(7648):105–109. doi:10.1038/nature21706

6. van Niel G, D’Angelo G, Raposo G. Shedding light on the cell biology of extracellular vesicles. Nat Rev Mol Cell Biol. 2018;19(4):213–228. doi:10.1038/nrm.2017.125

7. Raposo G, Stoorvogel W. Extracellular vesicles: exosomes, microvesicles, and friends. J Cell Biol. 2013;200(4):373–383. doi:10.1083/jcb.201211138

8. Kao CY, Papoutsakis ET. Extracellular vesicles: exosomes, microparticles, their parts, and their targets to enable their biomanufacturing and clinical applications. Curr Opin Biotechnol. 2019;60:89–98. doi:10.1016/j.copbio.2019.01.005

9. Thery C, Witwer KW, Aikawa E, et al. Minimal information for studies of extracellular vesicles 2018 (MISEV2018): a position statement of the International Society for Extracellular Vesicles and update of the MISEV2014 guidelines. J Extracell Vesicles. 2018;7(1):1535750. doi:10.1080/20013078.2018.1535750

10. Mulcahy LA, Pink RC, Carter DR. Routes and mechanisms of extracellular vesicle uptake. J Extracell Vesicles. 2014;3:24641. doi:10.3402/jev.v3.24641

11. Jiang J, Kao CY, Papoutsakis ET. How do megakaryocytic microparticles target and deliver cargo to alter the fate of hematopoietic stem cells? J Control Release. 2017;247:1–18. doi:10.1016/j.jconrel.2016.12.021

12. Wiklander OPB, Brennan MA, Lotvall J, Breakefield XO, El Andaloussi S. Advances in therapeutic applications of extracellular vesicles. Sci Transl Med. 2019;11(492):eaav8521. doi:10.1126/scitranslmed.aav8521

13. Jiang J, Woulfe DS, Papoutsakis ET. Shear enhances thrombopoiesis and formation of microparticles that induce megakaryocytic differentiation of stem cells. Blood. 2014;124(13):2094–2103. doi:10.1182/blood-2014-01-547927

14. Escobar C, Kao CY, Das S, Papoutsakis ET. Human megakaryocytic microparticles induce de novo platelet biogenesis in a wild-type murine model. Blood Adv. 2020;4(5):804–814. doi:10.1182/bloodadvances.2019000753

15. Wang W, Zuo B, Wang Y, et al. Megakaryocyte- and Platelet-Derived Microparticles as Novel Diagnostic and Prognostic Biomarkers for Immune Thrombocytopenia. J Clin Med. 2022;11(22):6776. doi:10.3390/jcm11226776

16. Stubbs JR, Homer MJ, Silverman T, Cap AP. The current state of the platelet supply in the US and proposed options to decrease the risk of critical shortages. Transfusion. 2021;61(1):303–312. doi:10.1111/trf.16140

17. Klein HG, Hrouda JC, Epstein JS. Crisis in the Sustainability of the U.S. Blood System. N Engl J Med. 2018;378(3):305–306. doi:10.1056/NEJMc1714807

18. Kao CY, Papoutsakis ET. Engineering human megakaryocytic microparticles for targeted delivery of nucleic acids to hematopoietic stem and progenitor cells. Sci Adv. 2018;4(11):eaau6762. doi:10.1126/sciadv.aau6762

19. Dellar ER, Hill C, Melling GE, Carter DRF, Baena-Lopez LA. Unpacking extracellular vesicles: RNA cargo loading and function. Journal of Extracellular Biology. 2022;1(5):e40. doi:10.1002/jex2.40

20. Kao CY, Jiang J, Thompson W, Papoutsakis ET. miR-486-5p and miR-22-3p Enable Megakaryocytic Differentiation of Hematopoietic Stem and Progenitor Cells without Thrombopoietin. Int J Mol Sci. 2022;23(10):5355. doi:10.3390/ijms23105355

21. Chattapadhyaya S, Haldar S, Banerjee S. Microvesicles promote megakaryopoiesis by regulating DNA methyltransferase and methylation of Notch1 promoter. Journal of Cellular Physiology. 2020;235(3):2619–2630. doi:10.1002/jcp.29166

22. Patel DB, Santoro M, Born LJ, Fisher JP, Jay SM. Towards rationally designed biomanufacturing of therapeutic extracellular vesicles: impact of the bioproduction microenvironment. Biotechnol Adv. 2018;36(8):2051–2059. doi:10.1016/j.biotechadv.2018.09.001

23. Ng CY, Kee LT, Al-Masawa ME, et al. Scalable Production of Extracellular Vesicles and Its Therapeutic Values: A Review. Int J Mol Sci. 2022;23(14):7986. doi:10.3390/ijms23147986

24. Syromiatnikova V, Prokopeva A, Gomzikova M. Methods of the Large-Scale Production of Extracellular Vesicles. Int J Mol Sci. 2022;23(18). doi:10.3390/ijms231810522

25. Hossler P, Khattak SF, Li ZJ. Optimal and consistent protein glycosylation in mammalian cell culture. Glycobiology. 2009;19(9):936–949. doi:10.1093/glycob/cwp079

26. Dunois-Larde C, Capron C, Fichelson S, Bauer T, Cramer-Borde E, Baruch D. Exposure of human megakaryocytes to high shear rates accelerates platelet production. Blood. 2009;114(9):1875–1883. doi:10.1182/blood-2009-03-209205

27. Ito Y, Nakamura S, Sugimoto N, et al. Turbulence Activates Platelet Biogenesis to Enable Clinical Scale Ex Vivo Production. Cell. 2018;174(3):636–648 e618. doi:10.1016/j.cell.2018.06.011

28. Luff SA, Papoutsakis ET. Megakaryocytic Maturation in Response to Shear Flow Is Mediated by the Activator Protein 1 (AP-1) Transcription Factor via Mitogen-activated Protein Kinase (MAPK) Mechanotransduction. J Biol Chem. 2016;291(15):7831–7843. doi:10.1074/jbc.M115.707174

29. Paganini C, Capasso Palmiero U, Pocsfalvi G, Touzet N, Bongiovanni A, Arosio P. Scalable Production and Isolation of Extracellular Vesicles: Available Sources and Lessons from Current Industrial Bioprocesses. Biotechnol J. 2019;14(10):e1800528. doi:10.1002/biot.201800528

30. Sumpio B, Chin J. Vessel Wall Biology. In: Cronenwett JL, Johnston KW, eds. Rutherford’s Vascular Surgery. 8 ed. Philadelphia: Saunders (Elsevier); 2014:34–48.

31. Bartolo MA, Qureshi MU, Colebank MJ, Chesler NC, Olufsen MS. Numerical predictions of shear stress and cyclic stretch in pulmonary hypertension due to left heart failure. Biomech Model Mechanobiol. 2022;21(1):363–381. doi:10.1007/s10237-021-01538-1

32. George SK, Laukova L, Weiss R, et al. Comparative Analysis of Platelet-Derived Extracellular Vesicles Using Flow Cytometry and Nanoparticle Tracking Analysis. Int J Mol Sci. 2021;22(8):3839. doi:10.3390/ijms22083839

33. Livak KJ, Schmittgen TD. Analysis of relative gene expression data using real-time quantitative PCR and the 2(-Delta Delta C(T)) Method. Methods. 2001;25(4):402–408. doi:10.1006/meth.2001.1262

34. Kondratov K, Nikitin Y, Fedorov A, et al. Heterogeneity of the nucleic acid repertoire of plasma extracellular vesicles demonstrated using high-sensitivity fluorescence-activated sorting. J Extracell Vesicles. 2020;9(1):1743139. doi:10.1080/20013078.2020.1743139

35. Perge P, Butz H, Pezzani R, et al. Evaluation and diagnostic potential of circulating extracellular vesicle-associated microRNAs in adrenocortical tumors. Sci Rep. 2017;7(1):5474. doi:10.1038/s41598-017-05777-0

36. Anand S, Samuel M, Kumar S, Mathivanan S. Ticket to a bubble ride: Cargo sorting into exosomes and extracellular vesicles. Biochim Biophys Acta Proteins Proteom. 2019;1867(12):140203. doi:10.1016/j.bbapap.2019.02.005

37. Chen Y, Zhao Y, Yin Y, Jia X, Mao L. Mechanism of cargo sorting into small extracellular vesicles. Bioengineered. 2021;12(1):8186–8201. doi:10.1080/21655979.2021.1977767

38. Leidal AM, Debnath J. Unraveling the mechanisms that specify molecules for secretion in extracellular vesicles. Methods. 2020;177:15–26. doi:10.1016/j.ymeth.2020.01.008

39. Villarroya-Beltri C, Baixauli F, Gutierrez-Vazquez C, Sanchez-Madrid F, Mittelbrunn M. Sorting it out: regulation of exosome loading. Semin Cancer Biol. 2014;28:3–13. doi:10.1016/j.semcancer.2014.04.009

40. Fabbiano F, Corsi J, Gurrieri E, Trevisan C, Notarangelo M, D’Agostino VG. RNA packaging into extracellular vesicles: An orchestra of RNA-binding proteins? J Extracell Vesicles. 2020;10(2):e12043. doi:10.1002/jev2.12043

41. Corrado C, Barreca MM, Zichittella C, Alessandro R, Conigliaro A. Molecular Mediators of RNA Loading into Extracellular Vesicles. Cells. 2021;10(12):3355. doi:10.3390/cells10123355

42. Chung J, Kim KH, Yu N, An SH, Lee S, Kwon K. Fluid Shear Stress Regulates the Landscape of microRNAs in Endothelial Cell-Derived Small Extracellular Vesicles and Modulates the Function of Endothelial Cells. Int J Mol Sci. 2022;23(3):1314. doi:10.3390/ijms23031314

43. Hergenreider E, Heydt S, Treguer K, et al. Atheroprotective communication between endothelial cells and smooth muscle cells through miRNAs. Nat Cell Biol. 2012;14(3):249–256. doi:10.1038/ncb2441

44. Xie F, Wen G, Sun W, et al. Mechanical stress promotes angiogenesis through fibroblast exosomes. Biochem Biophys Res Commun. 2020;533(3):346–353. doi:10.1016/j.bbrc.2020.04.159

45. Takafuji Y, Tatsumi K, Kawao N, Okada K, Muratani M, Kaji H. Effects of fluid flow shear stress to mouse muscle cells on the bone actions of muscle cell-derived extracellular vesicless. PLoS One. 2021;16(5):e0250741. doi:10.1371/journal.pone.0250741

46. Vechetti IJ, Jr., Peck BD, Wen Y, et al. Mechanical overload-induced muscle-derived extracellular vesicles promote adipose tissue lipolysis. FASEB J. 2021;35(6):e21644. doi:10.1096/fj.202100242R

47. Lv PY, Gao PF, Tian GJ, et al. Osteocyte-derived exosomes induced by mechanical strain promote human periodontal ligament stem cell proliferation and osteogenic differentiation via the miR-181b-5p/PTEN/AKT signaling pathway. Stem Cell Res Ther. 2020;11(1):295. doi:10.1186/s13287-020-01815-3

48. Wang YUE, Zheng Y, Li W. Exosomes derived from osteoclasts under compression stress inhibit osteoblast differentiation. Biocell. 2021;45(2):427–444. doi:10.32604/biocell.2021.013960

49. Xia B, Gao J, Li S, et al. Mechanical stimulation of Schwann cells promote peripheral nerve regeneration via extracellular vesicle-mediated transfer of microRNA 23b-3p. Theranostics. 2020;10(20):8974–8995. doi:10.7150/thno.44912

50. Najrana T, Mahadeo A, Abu-Eid R, et al. Mechanical stretch regulates the expression of specific miRNA in extracellular vesicles released from lung epithelial cells. J Cell Physiol. 2020;235(11):8210–8223. doi:10.1002/jcp.29476

51. Wang Y, Xie W, Feng Y, et al. Epithelial derived exosomes promote M2 macrophage polarization via Notch2/SOCS1 during mechanical ventilation. Int J Mol Med. 2022;50(1):96. doi:10.3892/ijmm.2022.5152

52. Yu W, Su X, Li M, et al. Three-dimensional mechanical microenvironment enhanced osteogenic activity of mesenchymal stem cells-derived exosomes. Chemical Engineering Journal. 2021;417:128040. doi:10.1016/j.cej.2020.128040

53. Flaumenhaft R, Dilks JR, Richardson J, et al. Megakaryocyte-derived microparticles: direct visualization and distinction from platelet-derived microparticles. Blood. 2009;113(5):1112–1121. doi:10.1182/blood-2008-06-163832

54. Das S, Harris JC, Winter EJ, Kao CY, Day ES, Papoutsakis ET. Megakaryocyte membrane-wrapped nanoparticles for targeted cargo delivery to hematopoietic stem and progenitor cells. Bioengineering & Translational Medicine. 2022:e10456. doi:10.1002/btm2.10456

55. Vion AC, Ramkhelawon B, Loyer X, et al. Shear stress regulates endothelial microparticle release. Circ Res. 2013;112(10):1323–1333. doi:10.1161/CIRCRESAHA.112.300818

56. Boulestreau J, Maumus M, Rozier P, Jorgensen C, Noel D. Mesenchymal Stem Cell Derived Extracellular Vesicles in Aging. Front Cell Dev Biol. 2020;8:107. doi:10.3389/fcell.2020.00107

57. McVey MJ, Weidenfeld S, Maishan M, et al. Platelet extracellular vesicles mediate transfusion-related acute lung injury by imbalancing the sphingolipid rheostat. Blood. 2021;137(5):690–701. doi:10.1182/blood.2020005985

58. Belliveau J, Papoutsakis ET. Extracellular vesicles facilitate large-scale dynamic exchange of proteins and RNA among cultured Chinese hamster ovary and human cells. Biotechnol Bioeng. 2022;119(5):1222–1238. doi:10.1002/bit.28053

59. Wei Z, Batagov AO, Schinelli S, et al. Coding and noncoding landscape of extracellular RNA released by human glioma stem cells. Nat Commun. 2017;8(1):1145. doi:10.1038/s41467-017-01196-x

60. Chevillet JR, Kang Q, Ruf IK, et al. Quantitative and stoichiometric analysis of the microRNA content of exosomes. Proc Natl Acad Sci U S A. 2014;111(41):14888–14893. doi:10.1073/pnas.1408301111

61. Stevanato L, Thanabalasundaram L, Vysokov N, Sinden JD. Investigation of Content, Stoichiometry and Transfer of miRNA from Human Neural Stem Cell Line Derived Exosomes. PLoS One. 2016;11(1):e0146353. doi:10.1371/journal.pone.0146353

62. Vion AC, Birukova AA, Boulanger CM, Birukov KG. Mechanical forces stimulate endothelial microparticle generation via caspase-dependent apoptosis-independent mechanism. Pulm Circ. 2013;3(1):95–99. doi:10.4103/2045-8932.109921

63. Cunin P, Penke LR, Thon JN, et al. Megakaryocytes compensate for Kit insufficiency in murine arthritis. J Clin Invest. 2017;127(5):1714–1724. doi:10.1172/JCI84598

64. Fuhrken PG, Apostolidis PA, Lindsey S, Miller WM, Papoutsakis ET. Tumor suppressor protein p53 regulates megakaryocytic polyploidization and apoptosis. J Biol Chem. 2008;283(23):15589–15600. doi:10.1074/jbc.M801923200

