## Supplemental Data for "Similar but distinct: the impact of biomechanical forces and culture age on the production, cargo loading, and biological efficacy of human megakaryocytic extracellular vesicles for applications in cell and gene therapies"

**This document includes:**

**Figure S1:**	Total miRNA content for MkEVs from the shake flask experiments.

**Figure S2:**	Total miRNA content for MkEVs and parent cells from the syringe pump experiments.

**Figure S3:**	Total and individual miRNA content in MkEVs from different days.

**Figure S4:**	Bioactivity of MkEVs produced on different days.

**Figure S1**

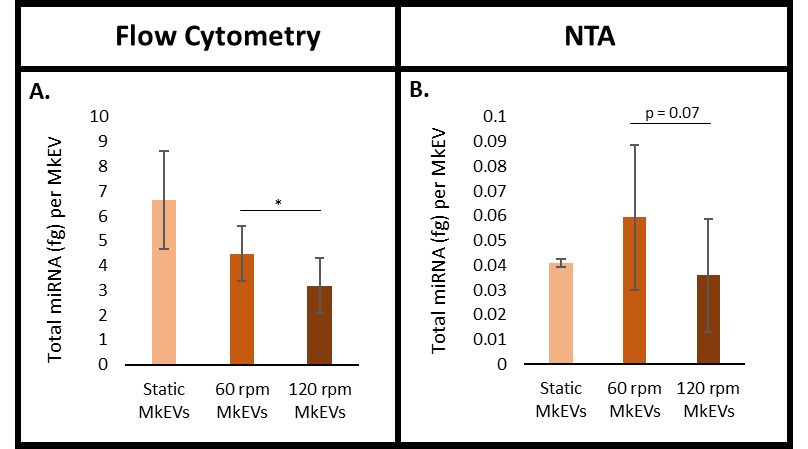

**Figure S1. Total miRNA content for MkEVs from the shake flask experiments.** (A) Total miRNA (in femtograms) per MkEV for flow cytometry-based MkEV counts. (B) Total miRNA (in femtograms) per MkEV for NTA-based MkEV counts. Error bars indicate SEM of 3 biological replicates. Paired Student’s t-tests were performed on all data; * represents p < 0.05.

**Figure S2**

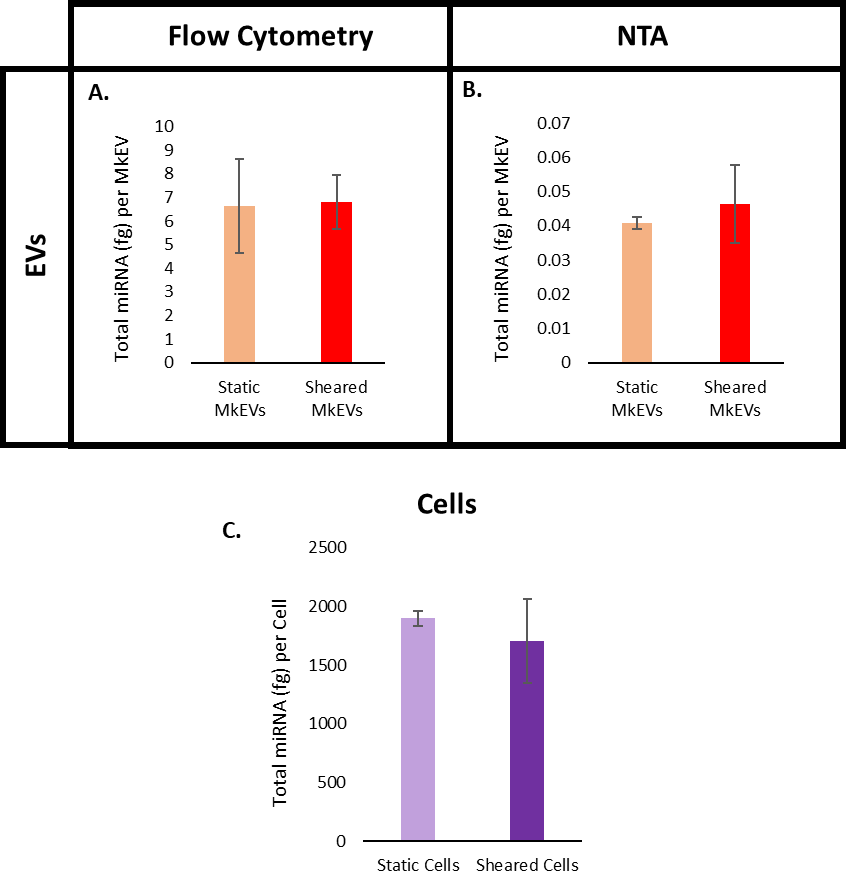

**Figure S2. Total miRNA content for MkEVs and parent cells from the syringe pump experiments.** (A) Total miRNA (in femtograms) per MkEV for flow cytometry-based MkEV counts. (B) Total miRNA (in femtograms) per MkEV for NTA-based MkEV counts. (C) Total miRNA per Mk following syringe pump-induced shear or control treatment. Error bars indicate SEM of 3 biological replicates. Unpaired Student’s t-tests were performed on all data.

**Figure S3**

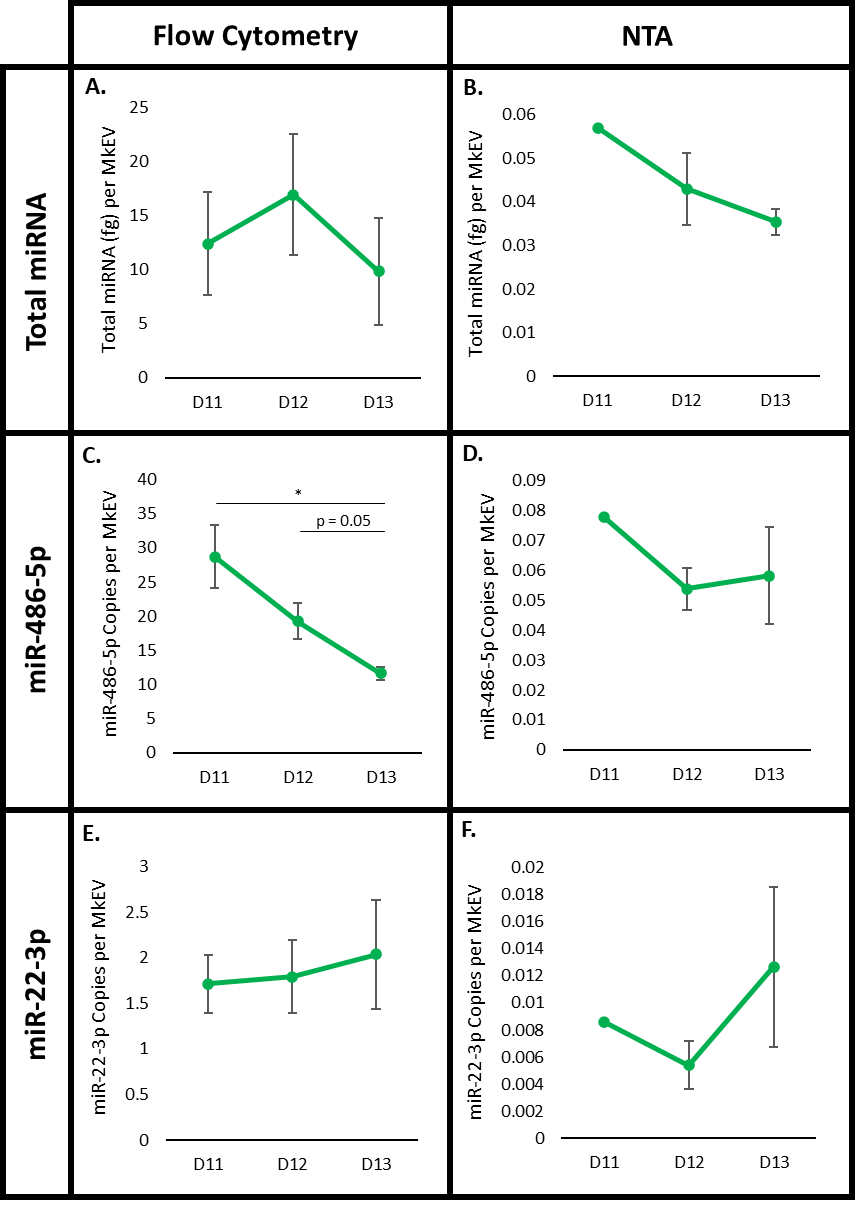

**Figure S3. Total and individual miRNA content in MkEVs from different days.** (A) Total miRNA (in femtograms) per MkEV from D11-D13 for flow cytometry-based MkEV counts. (B) Total miRNA per MkEV from D11-D13 for NTA-based MkEV counts. (C) Copies of miR-486-5p per MkEV from D11-D13 for flow cytometry-based MkEV counts. (D) Copies of miR-486-5p per MkEV from D11-D13 for NTA-based MkEV counts. (E) Copies of miR-22-3p per MkEV from D11-D13 for flow cytometry-based MkEV counts. (F) Copies of miR-22-3p per MkEV from D11-D13 for NTA-based MkEV counts. Error bars indicate SEM of 3 biological replicates; data points using NTA-derived D11 MkEV counts consist of 1 biological replicate each. Paired Student’s t-tests were performed on all data; * represents p < 0.05.

**Figure S4**

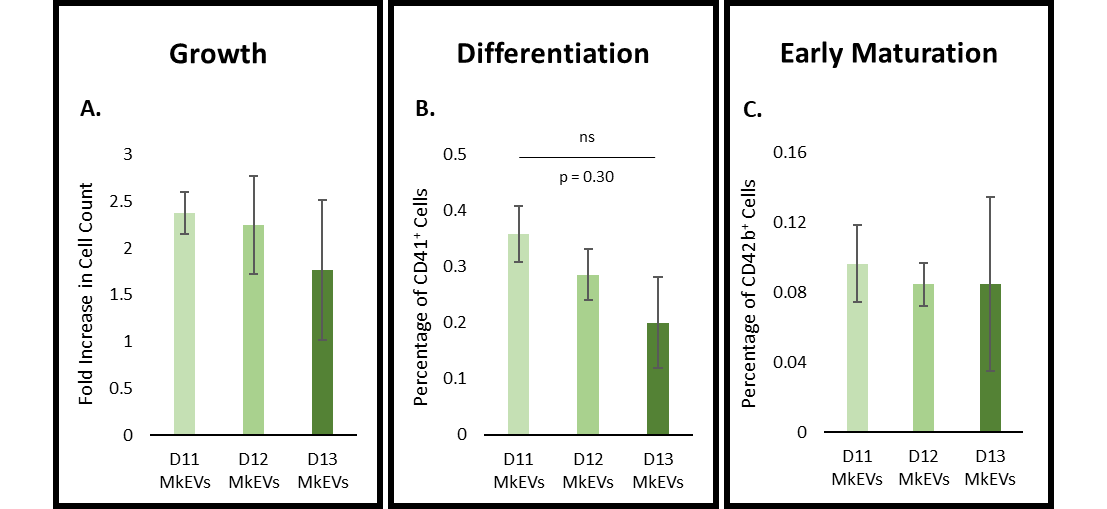

**Figure S4. Bioactivity of MkEVs produced on different days.** MkEVs from D11, D12, and D13 Mk cultures were co-cultured with HSPCs at a 20:1 ratio for 7 days. (A) Fold change in cell growth (relative to untreated cells) following co-culture with various MkEV samples. (B) The percentage of cells in each co-culture expressing CD41 (an Mk marker). (C) The percentage of cells in each co-culture expressing CD42b (a marker for early Mk maturation). Error bars indicate SEM of 3 biological replicates, except for (A), where one outlier was omitted (leaving 2 biological replicates). Paired Student’s t-tests were performed on all data; ns represents non-significance.
